# Unraveling the Co-Morbidity between COVID-19 and Neurodegenerative Diseases Through Multi-scale Graph Analysis: A Systematic Investigation of Biological Databases and Text Mining

**DOI:** 10.1101/2025.02.13.637069

**Authors:** Negin Sadat Babaiha, Stefan Geissler, Vincent Nibart, Heval Atas Güvenilir, Vinay Srinivas Bharadhwaj, Alpha Tom Kodamullil, Juergen Klein, Marc Jacobs, Martin Hofmann-Apitius

## Abstract

The COVID-19 pandemic has generated a vast volume of research, yet much of it focuses on individual diseases, overlooking complex comorbidity relationships. While extensive literature exists on both neurodegenerative diseases (NDDs), such as Alzheimer’s and Parkinson’s, and COVID-19, their intersection remains underexplored. Co-morbidity modeling is crucial, particularly for hospitalized patients often presenting with multiple conditions.

This study investigates the interplay between COVID-19 and NDDs by integrating knowledge graphs (KGs) built from curated biomedical datasets and text mining tools. We performed comprehensive analyses—including path analysis, phenotype coverage, and mapping of cellular and genetic factors—across multiple KGs, such as PrimeKG, DrugBank, OpenTargets, and those generated via natural language processing (NLP) methods.

Our findings reveal notable variability in graph density and connectivity, with each KG offering unique insights into molecular and phenotypic links between COVID-19 and NDDs. Key genetic and inflammatory markers, especially immune response genes, consistently appeared across graphs, suggesting a shared pathogenic basis.

By unifying structured biological data with unstructured textual evidence, we enhance co-morbidity modeling and improve recall in identifying mechanisms underlying COVID-19– NDD interactions. This integrative framework supports the development of a co-morbidity hypothesis database aimed at facilitating therapeutic target discovery.

All data, methods, and instructions for accessing the co-morbidity hypothesis database are publicly available at: https://github.com/SCAI-BIO/covid-NDD-comorbidity-NLP.

## 1 Introduction

The COVID-19 pandemic, which emerged in late 2019, was swiftly declared an international health emergency by the World Health Organization (WHO) due to its rapid global spread and severe impact on public health [1], [2], [3]. The disease, caused by the severe acute respiratory syndrome coronavirus 2 (SARS-CoV-2), resulted in severe mortality and emergency cases worldwide. The COVID-19 pandemic has posed a complex and multifaceted challenge, overwhelming global healthcare systems with an unprecedented surge in patient numbers and severe resource shortages, while also triggering widespread social and economic disruptions across the globe [4], [5]. Besides, the pervasive uncertainty surrounding the virus and its effects has exacerbated mental health issues, with a marked increase in cases of depression, anxiety, and burnout among the global population [1], [6].

COVID-19 disease primarily targets the respiratory system, but it has also been associated with a wide range of symptoms and complications affecting multiple organ systems such as the cardiovascular and immune system [1], [7]. Recent studies have revealed complex immune dysregulation in COVID-19, particularly in severe cases [8]. Macrophages and monocytes, key components of the innate immune system, play critical roles in pathogen recognition and inflammation control [9]. Macrophages, as tissue-resident cells, initiate local immune responses, while monocytes are recruited from blood during infection. The disease progression follows a time-dependent model where initial interferon response is crucial for viral control [8]. However, delayed or prolonged responses can trigger hyperinflammation, characterized by excessive mononuclear phagocyte activation and dysregulated tissue repair. Severe cases exhibit distinct immunological changes, including elevated pro-inflammatory cytokines, neutrophilia, lymphopenia, and reduced HLA-DR expression on monocytes [8].

Beyond its well-documented respiratory manifestations and immune dysregulation, from the start of the pandemic, the disease has been linked to a range of neurological complications [10]. Early in the pandemic, reports began to surface detailing these complications, which varied from mild symptoms such as headaches and myalgia [11], to more severe and potentially life-threatening conditions, including seizures, strokes, and Guillain-Barré syndrome [12]. Subsequent in vitro studies indicated the potential for SARS-CoV-2 to infect neurons and astrocytes, raising concerns about direct viral involvement in neurological dysfunction [13]. However, findings from autopsy studies suggest that such dysfunction is unlikely to result from direct viral invasion in vivo [12], [14]. Instead, these studies propose that the virus affects the brain indirectly, potentially through mechanisms such as immune cell activation, the release of peripherally generated inflammatory mediators, or alterations in the blood-brain barrier [12], [13], [14], [15]. Nonetheless, preliminary evidence hints towards a direct aggregation potential for soluble amyloid beta of SARS-CoV-2, and HIV, as highlighted in a recent preprint [16]. In their study, they demonstrate that certain viruses, including SARS-CoV-2 and HSV-1, can induce the aggregation of proteins involved in neurodegenerative diseases (NDDs) through a physicochemical process known as heterogeneous nucleation (HEN). This mechanism allows viruses to catalyze protein clumping in cerebrospinal fluid (CSF) without requiring viral replication, suggesting that these viral infections may directly contribute to the development of NDDs by triggering protein aggregation pathways [16]. Recent comprehensive analyses have further elucidated the complex relationship between COVID-19 and NDDs [17]. More than two-thirds of hospitalized COVID-19 patients experience incomplete recovery even months after infection, with specific implications for neurological health. As mentioned in their study, the virus’ neurological impact appears to operate through five distinct pathways: the olfactory epithelial route, blood-brain barrier penetration, lateral ventricles and choroid plexus involvement, vagus nerve transmission, and corneal epithelial pathway [17]. These multiple entry routes may explain some of the viruś diverse neurological effects. Particularly concerning is the virus’s interaction with existing neurodegenerative conditions: Alzheimer’s disease patients show increased COVID-19 susceptibility and mortality risk, while Parkinson’s disease patients demonstrate a 58% higher hospitalization rate compared to healthy controls [17]. In multiple sclerosis patients, approximately 29% experience prolonged COVID-19 symptoms lasting over four weeks, predominantly manifesting as fatigue [17]. The emergence of sophisticated diagnostic approaches, including FDG-PET imaging and EEG monitoring, has enabled better assessment of these neurological impacts. While direct causal relationships between SARS-CoV-2 and NDDs remain under investigation, mounting evidence suggests that COVID-19 may act as both a trigger for new neurodegenerative conditions and an accelerator of existing ones, particularly through mechanisms of sustained inflammation and immune response dysregulation [17].

Given these findings, there is a growing interest in understanding the relationship between COVID-19 and the development of NDDs, such as Alzheimer’s disease (AD). To investigate the mechanisms by which COVID-19 may cause neurological complications, a research group conducted a study focused on the neurological aspects of COVID-19 under a project known as CIAO (https://www.ciao-covid.net). This project aimed to uncover the existing knowledge on COVID-19 pathogenesis by utilizing the Adverse Outcome Pathway (AOP) framework [18]. The AOP framework systematically maps disease progression from an initial molecular event, such as SARS-CoV-2 infection, through key biological processes, ultimately leading to adverse outcomes like respiratory distress, organ failure, insomnia, or cognitive impairments such as brain fog [19]. However, the rapidly evolving variants of COVID-19 and the broad spectrum of clinical symptoms present significant challenges to applying this framework effectively. Given the vast amount of research, clinical trials, and data generated in the years following the COVID-19 pandemic, there is a wealth of valuable information that can be leveraged to better understand the underlying mechanisms. The large volume of unstructured textual data makes natural language processing (NLP) and text mining particularly valuable tools in this endeavor. For instance, a recent study employed NLP and explored the corpora around neurological toxicity and COVID-19 by developing a language model named NeuroCORD [20]. This model leverages Bidirectional Encoder Representations from Transformers (BERT) embeddings and has been trained on a corpus of literature abstracts to classify and distinguish COVID-19 articles that focus on neurological disorders from those covering other topics [20]. In the contentious debate over COVID-19 vaccinations, a widely cited study used text mining on Twitter (now “X”) data to assess public hesitancy, offering key insights into societal responses to the pandemic [21]. Building on this approach, other researchers have employed techniques like co-occurrence analysis, clustering, and topic modeling, to investigate the common manifestations of COVID-19 and identify potential therapeutic agents [22].

Despite the vast amount of research on NDDs such as AD and Parkinson’s (PD), and the rapidly expanding body of literature on COVID-19, there remains a significant gap when it comes to studying the crosstalk between these two conditions. The dominance of “one-disease-centric” research presents a challenge for mining information on comorbidities, which are a critical aspect of real-world health dynamics. In clinical settings, patients rarely present with a single disease; comorbidities are the norm, particularly in those hospitalized. Early studies of hospitalized COVID-19 patients, conducted through the ISARIC4C protocol [23], revealed that over three-quarters had at least one co-morbidity, with conditions such as cardiac disease, pulmonary disease, chronic kidney disease, obesity, cancer, neurological disorders, dementia, and liver disease all associated with higher in-hospital mortality [24]. Analysis of primary care data from 40% of the English population further revealed hypertension (34.3%), asthma (15.9%), and diabetes (9.9%) as the most common comorbidities [24]. More recent research has shifted focus to multimorbidity and revealed that among 1,706 severe COVID-19 cases in the UK Biobank [25], 25.3% of patients had multiple conditions, with stroke and hypertension being the most prevalent combination, and chronic kidney disease paired with diabetes showing the highest associated risk (OR 4.93; 95% CI 3.36-7.22) [24]. In a separate study, Romagnolo et al. investigated the influence of pre-existing neurological conditions on COVID-19 outcomes in a cohort of 332 patients, of whom 22.6% were diagnosed with neurological disorders [25]. Their study found that patients with neurological diseases had a significantly higher case fatality rate (48.0% vs. 24.0%) and were independently associated with increased mortality. Specifically, NDDs were linked to the highest mortality (73.9% vs. 39.1%), while cerebrovascular diseases showed a higher, though non-significant, mortality rate. These patients were also older, had more comorbidities, and presented with more severe COVID-19 symptoms. The unprecedented surge in studies regarding COVID-19-related comorbidities, although offering critical insights, often includes speculation and frequently falls short of delivering the robust, evidence-based data necessary for drawing definitive conclusions about COVID-19 comorbidities. Most of these insights are derived from large studies that focus only on hospitalized patients, which introduces bias into the findings [24]. Additionally, in general population studies, factors such as non-random sampling may distort the results, further complicating the interpretation of the data [24], [26]. Besides, robust insights into comorbidities between COVID-19 and NDDs primarily come from observational cohort studies, such as UK Biobank [27], or from experimental systems like organoid studies and blood-brain barrier (BBB) models, but these approaches can limit the generalizability of the findings.

Our current study, conducted within the framework of the COMMUTE project (www.commute-project.eu), aims to bridge existing gaps by integrating knowledge from both neurological diseases and COVID-19 research. Utilizing a graph-based approach, we model and mine co-morbidity relationships to gain a deeper understanding of their interactions. This methodology enables us to synthesize a vast and fragmented body of literature into a unified framework, facilitating a comprehensive analysis of existing knowledge stored in databases through the lens of Knowledge Graphs (KGs). By integrating diverse data sources, our approach not only enhances the understanding of the relationships between SARS-CoV-2 infection and NDDs but also offers novel insights into how COVID-19 may elevate the risk of NDDs at both the population and individual levels. The graph-based approach provides several advantages, such as visualizing complex relationships, integrating heterogeneous data sources, identifying hidden patterns, and allowing for predictive modeling of potential co-morbidities. KGs are structured data formats designed to represent, organize, and analyze complex biological and medical information by capturing intricate relationships between diverse entities. Constructing disease-specific co-morbidity KGs can be approached from several perspectives, including manual curation, database integration, and the use of automated tools. Integrating curated databases provides a robust and reliable method for creating these KGs, especially for complex diseases like COVID-19. Curated databases offer high-quality, validated data, ensuring the accuracy and completeness of the resulting graphs, which enhances their effectiveness in analyzing and understanding disease mechanisms.

For instance, DisGeNET [28] is a widely used database that catalogs disease-gene and disease-variant associations, aggregating information from trusted sources like UniProt [29], ClinVar [30], and the GWAS Catalog [31]. Integrating curated data from such resources facilitates the creation of detailed, precise disease-gene relationship maps. Databases such as OpenTargets [32], DrugBank [33], and PrimeKG [34] further enrich the depth and utility of KGs. However, there are limitations to this approach. Relying solely on curated resources can restrict the scope of the KG, as these databases may lag in capturing the latest research findings and emerging trends still under investigation. Moreover, the curation process is inherently subjective and may introduce bias, as it relies on the judgment of curators. Additionally, integrating data from multiple sources can be challenging due to variations in data formats, standards, and ontologies, and curated databases often contain redundant, inconsistent, or outdated records, complicating the integration process despite efforts to address these issues.

In contrast, constructing KGs through text mining of unstructured sources, such as scientific literature, offers a dynamic and complementary approach. Frameworks like the Integrated Network and Dynamical Reasoning Assembler (INDRA) (https://github.com/bgyori/indra) exemplify this method by automating the extraction and assembly of mechanistic models from textual data. Leveraging Natural Language Processing (NLP) techniques like REACH (https://indra.readthedocs.io/en/stable/modules/sources/reach/index.html) and TRIPS (https://indra.readthedocs.io/en/stable/modules/sources/trips/index.html), INDRA identifies entities such as genes and proteins and their interactions, structuring these interactions into standardized statements linked to database identifiers to ensure accuracy. This process allows for the discovery of novel associations that may not yet be captured by curated databases. While text mining offers flexibility and the potential to uncover new associations, it also presents challenges, including the need to continuously optimize and retrain NLP models to adapt to new datasets and terminologies. Furthermore, the inherent variability and ambiguity of natural language can introduce errors, highlighting the need for rigorous curation to maintain the quality and reliability of KGs.

Curated databases and text mining offer complementary strengths, and their integration holds great potential for advancing KG construction and biomedical discovery. In this study, we combine comprehensive database investigation and text mining to build mechanism-based co-morbidity KGs. By focusing on the mechanistic understanding of co-morbidity between COVID-19 and NDDs, we explore how different data sources and tools contribute to generating testable hypotheses. Our goal is to optimize co-morbidity modeling by leveraging the strengths of these resources, ultimately creating a comprehensive database of hypotheses that not only reflects current knowledge on co-morbidity mechanisms but also uncovers new insights derived from graph mining across the integrated data.

## 2 Materials and Methods

Our workflow starts with collecting co-morbidity-specific information on COVID-19 and NDDs, specifically AD and PD, in curated databases. This initial step provides a foundational understanding of potential associations and interactions, ensuring a data-driven approach to subsequent analyses. After collecting the data, the interactions are systematically integrated into KG representations, enabling structured visualization and analysis of the relationships within each data source. These data sources are often stored in diverse formats, necessitating transformation and preprocessing to ensure compatibility with the task at hand. For instance, some databases primarily focus on single relationship types, such as disease-gene associations, while others encompass multiple relationship types across various biological entities. To address these disparities, custom scripts are developed to handle data extraction and preprocessing, manage various files, and construct graphs that accurately reflect the underlying data structure. Once prepared, the processed graphs are uploaded to Neo4j graph database (https://neo4j.com), enabling efficient storage, querying, and analysis of the integrated data. This workflow ensures a cohesive and scalable approach to handling heterogeneous datasets and supports the creation of comprehensive KGs for advanced analyses.

In parallel to the information retrieval from databases, we constructed a co-morbidity KG through information extraction from relevant literature. This began with a search of PubMed for publications addressing COVID-19 and NDD correlations in recent years, as detailed in section 2.1. Once the textual data was retrieved, filtered, and the relevance of the selected publications was validated, NLP techniques were employed to identify key entities and extract their relationships. These techniques enabled the systematic analysis of complex biomedical concepts, facilitating the mapping of interactions and associations critical to understanding the correlations between COVID-19 and NDDs. The resulting KG captures a wide spectrum of entities, and their interactions related to the COVID-19 and NDD co-morbidity.

A comparative analysis was then conducted between the KGs derived from databases and those constructed via text mining. This analysis leveraged graph-based path analysis and reasoning algorithms to uncover central nodes, biomarkers, drug targets, and identifiable disease pathways. An overview of the workflow is shown in Figure 1.

**Figure 1.**
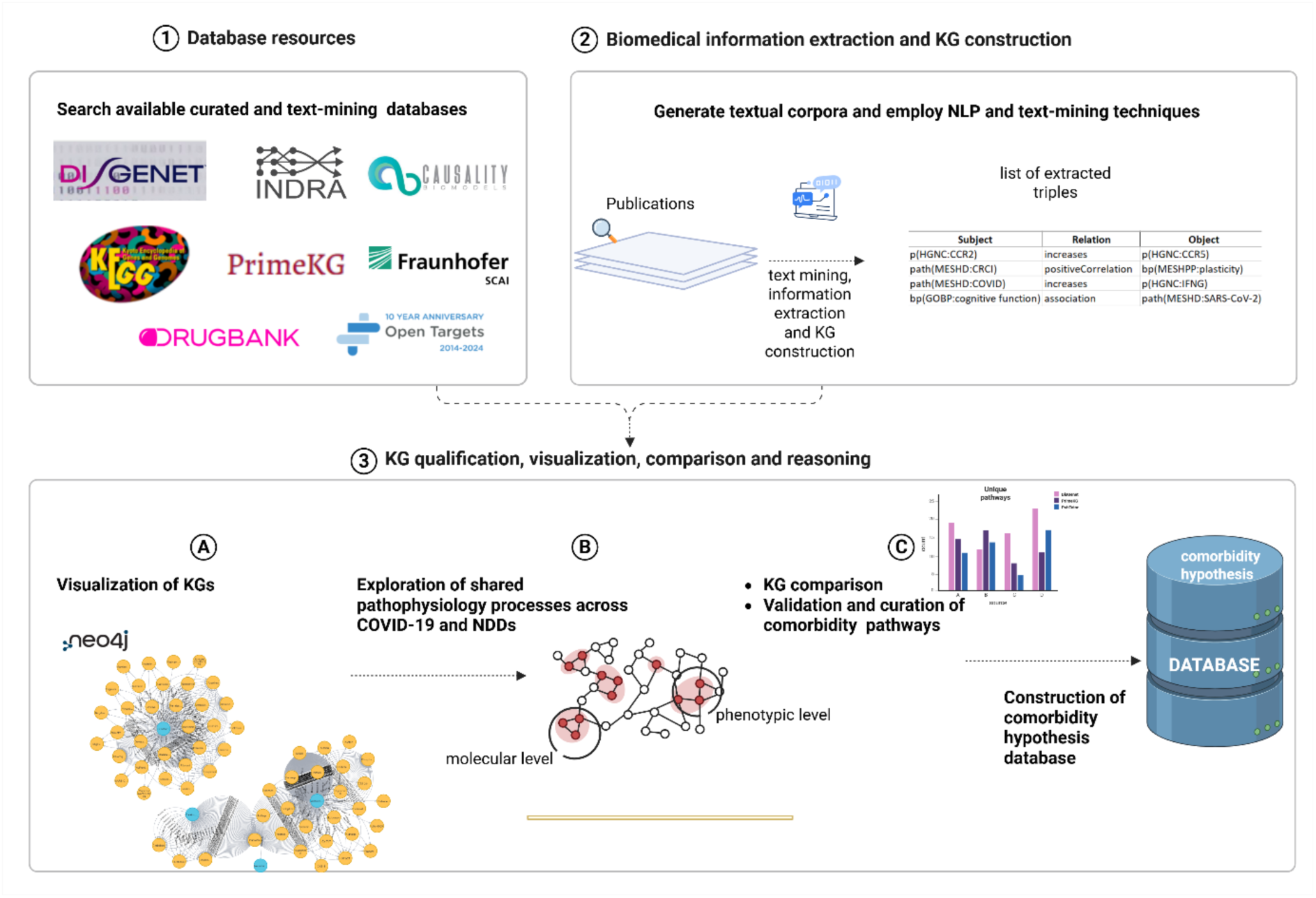
Overview of the co-morbidity KG construction and analysis pipeline. The workflow integrates data on COVID-19 and NDDs from curated databases (1) and PubMed publications to construct KGs (2). Using NLP techniques, relationships between key biomedical entities are extracted and visualized via Neo4j(2,3-A), facilitating the identification of critical nodes, biomarkers, and shared disease pathways (3-B). Comparative analysis of KGs from databases and text mining uncovers insights into the interactions between COVID-19 and NDDs (C). Finally, the candidate hypothesis reflecting potential co-morbidity between COVID-19 and NDDs is stored in a database, which will be further enriched and validated.

### 2.1 Textual Data

#### 2.1.1 PubMed Search

The overview of textual corpora generation and processing is represented in Figure 2. We started our corpora generation with a focused search in the PubMed database, using a combination of keywords and Medical Subject Headings (MeSH) (https://www.ncbi.nlm.nih.gov/mesh/) terms. We thoroughly reviewed recent relevant literature to find key topics and emerging trends in research on COVID-19 and its effects on the nervous system. We paid special attention to highly relevant and frequently cited papers to identify common keywords and phrases used by experts. We also consulted with specialists in neurology and infectious diseases to make sure our search strategy was accurate and relevant. As a result, we created a specific list of search terms to look for such as combinations like “COVID-19 AND Blood-Brain Barrier Disruption,” “COVID-19 AND Neuronal Infection,” and “COVID-19 AND Cerebrovascular System,” among others (a complete list of the search terms can be found in the Supplementary File, section 1.). We utilized the E-utility public API [35] to retrieve PMIDs for the relevant search term list. To ensure the inclusion of the most current research, we focused on articles published between 2021 and 2024.

**Figure 2.**
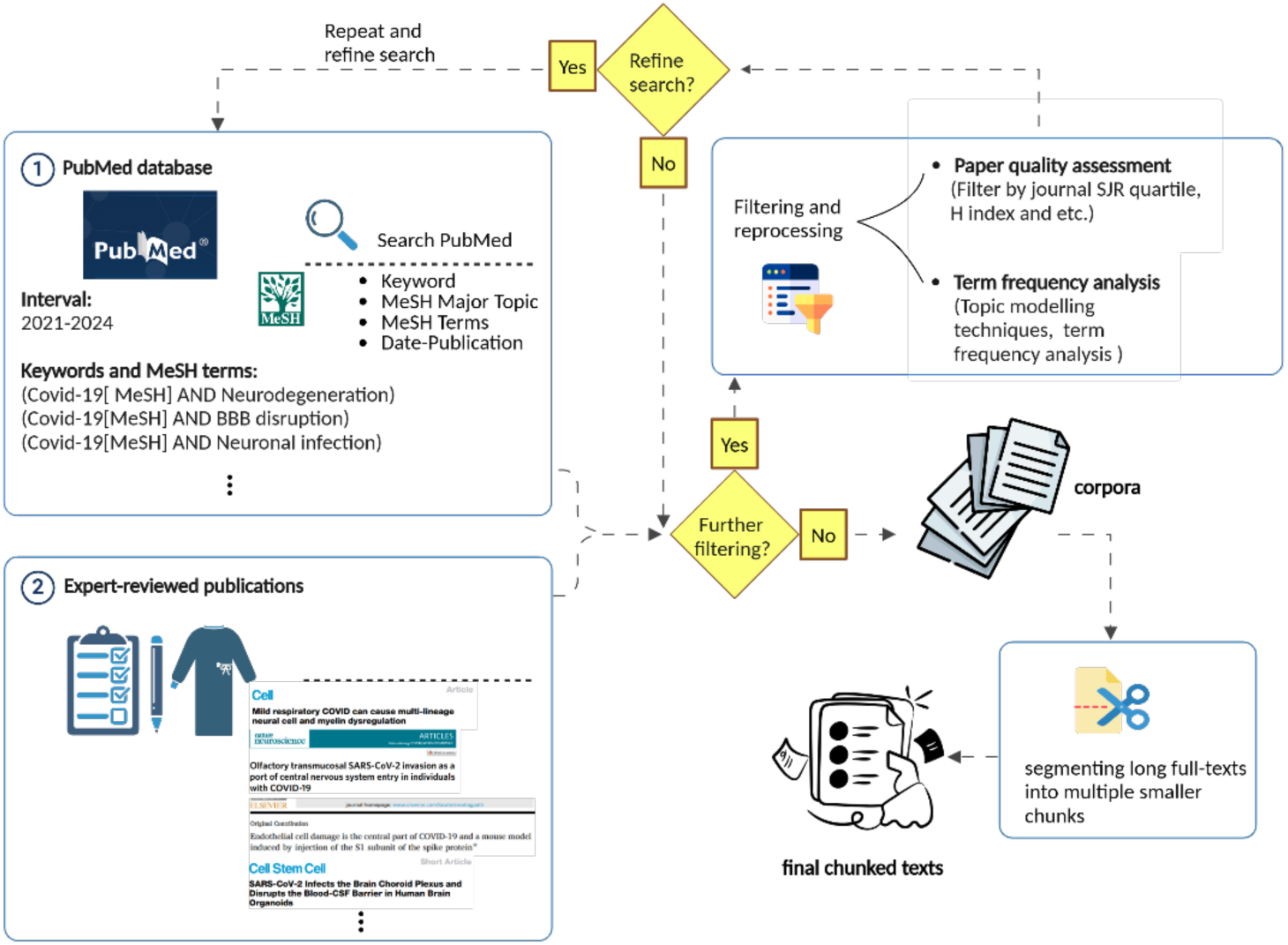
The strategies used to generate and refine a corpus of scientific literature focused on the co-morbidity between COVID-19 and NDDs. Initially, we conducted a search on PubMed using relevant keywords and MeSH terms related to COVID-19-NDD co-morbidity and defined appropriate time intervals. Consultation with domain experts allowed us to select a highly relevant set of corpora. The extracted literature underwent a post-processing and assessment phase, utilizing topic modeling and journal evaluations. As the search terms were refined and updated, the process became more comprehensive, involving an expanded search on PubMed. Finally, the list of selected publications was divided into manageable chunks, preparing them for further processing by NLP models.

#### 2.1.2 Expert Curation

In addition to conducting an extensive search using the E-utility API, we curated a targeted selection of papers based on expert recommendations. These recommendations were provided by specialists in Clinical Neurology and Infectious Diseases, who are part of the COMMUTE project consortium. This consortium represents leading research institutions from Germany, Spain, Luxembourg, and The Netherlands, ensuring that the selection process was informed by a diverse and highly qualified panel of experts. This combined approach allowed us to capture a comprehensive and focused set of publications relevant to the intersection of COVID-19 and NDDs. A complete list of the PMIDs of these publications is provided in our project GitHub repository data (https://github.com/SCAI-BIO/covid-NDD-comorbidity-NLP).

### 2.2 Textual Corpora Assessment

To enhance our PubMed search strategy and ensure the most relevant papers were included in our corpus, we reviewed term frequencies to identify any overlooked papers and important terms that were absent from our initial search. We analyzed how often our search terms appeared in the full text of the publications. We observed that some MeSH terms might have been missing because newly published articles had not yet been assigned these terms. We then incorporated these terms into our search, refining our strategy for a more comprehensive review. The corpora generated through keyword and MeSH term searches, however, may still contain irrelevant papers. To further filter the publications and identify the most related ones, we employed several strategies: 1) ranking publications and 2) employment of topic modeling. First, we filtered papers based on their publication ranking in high-quality journals within the domain. Additionally, we applied topic modeling and term frequency analysis to extract the most pertinent topics for each set of corpora. Details of these approaches are outlined in the Supplementary File, section 2.

### 2.3 Pre-Processing of Textual Corpora

To process the textual corpus with NLP tools, we employed the BioC API for PubMed Central (PMC) [36] to extract full-text documents. Of the 100 documents targeted, 91 were successfully retrieved and freely accessible. Given the large size of the full-text documents, we optimized our processing by segmenting each document into paragraphs and treating them separately. Each document was typically divided into 100 to 300 paragraphs. This method ensured efficient processing and manageable data handling for our NLP pipeline.

### 2.4 Co-morbidity KG Construction

The comprehensive collection of co-morbidity KGs was constructed using two main methods, as previously shown in Figure 1. First, we integrated data from trusted biomedical databases and curated sources to gather reliable information. Second, we used NLP and text mining to extract useful data directly from scientific texts. The following sections will explain each method in more detail.

#### 2.4.1 Available Databases and Sources

##### 2.4.1.1 KEGG Pathway Database

We utilized the KEGG Pathway database [37] to gather detailed information on COVID-19 and NDDs. Using the KEGG API, we retrieved relevant data for each disease by first extracting KEGG IDs for each COVID-19, AD, PD, and NDDs (general), and then obtaining comprehensive disease-specific information, including associated pathways, genes, and drugs. This provided valuable insights into the genetic, molecular, and therapeutic aspects of these diseases. The technical details for data extraction from KEGG are added to the Supplementary File, section 3.

##### 2.4.1.2 DisGeNET

DisGeNET, a database that collects over 380,000 gene-disease associations across more than 16,000 genes and 13,000 diseases, was used to gather key information on COVID-19, AD, PD, and NDDs. By leveraging the DisGeNET R package, we extracted Unified Medical Language System (UMLS) IDs for each disease and then used DisGeNET API to identify relevant disease-gene associations to build a KG for further analysis and comparison.

##### 2.4.1.3 DrugBank

DrugBank, a trusted resource offering comprehensive details on drugs, including chemical properties, mechanisms of action, and protein-gene interactions, was utilized in our study. We accessed DrugBank version 5.1.12, which includes 2,358 approved drugs and nearly 366,000 drug-drug interactions. This data was filtered to focus specifically on drugs associated with COVID-19, AD, PD, and NDDs for further analysis.

##### 2.4.1.4 OpenTargets

OpenTargets is a comprehensive platform that aggregates and refines data from various public sources, scoring and ranking target-disease associations based on genomics, transcriptomics, and other biological data. We utilized OpenTargets (v24.06) to identify agents targeting COVID-19 and NDDs by accessing the platform’s user interface and downloading targets for each disease individually for further analysis.

##### 2.4.1.5 INDRA Database

INDRA is a source developed at Harvard Medical School using automated extraction of molecular mechanisms from both text and curated databases. It compiles mechanistic knowledge into a machine-readable format called “statements” and integrates these into the non-redundant knowledge base. Using the INDRA API (v1.0), we extracted disease-related statements for both COVID and NDD and only considered the statements where the confidence score was higher than 0.85.

##### 2.4.1.6 PrimeKG

PrimeKG, introduced in 2023, is a comprehensive graph that integrates multi-dimensional disease-related data from 20 high-quality biomedical resources. It maps over 17,000 diseases with more than 4 million connections, encompassing disease-related proteins, biological processes, pathways, anatomical structures, and therapeutic compounds.

To investigate the relationship between COVID-19 and NDDs, we leveraged PrimeKG’s extensive KG by searching for both COVID-19 (e.g., SARS-CoV-2, COVID-19, coronavirus) and NDDs (e.g., Alzheimer’s, Parkinson’s, ALS). In total, there are x nodes and y edges in the COVID-NDD subgraph extracted from PrimeKG.

##### 2.4.1.7 Causality Biomodels (CBM)

CBM is a company (https://causalitybiomodels.com), specializing in bio-curation and knowledge extraction. [*For transparency reasons:* CBM is a spin-off started by and owned by members of Fraunhofer SCAI.] By extracting semantic information from published sources, CBM develops valuable knowledge models in the life sciences, a capability demonstrated in our previous two papers [38], [39]. The textual corpora discussed in section 2.1, which were reviewed by domain experts, were manually curated by CBM, resulting in the extraction of over 3,000 mechanism triples encoded in Biological Expression Language (BEL) (https://bel.bio). This curated graph is also used as a gold standard for subsequent comparisons.

##### 2.4.1.8 Manually Curated Disease Maps for COVID-19 and NDDs

We also utilized KGs developed by colleagues at the Fraunhofer Institute for Algorithms and Scientific Computing (SCAI) to analyze the co-morbidity between COVID-19 and NDDs. The COVID-19 KG, developed by Domingo-Fernández et al. in 2021, is a comprehensive cause-and-effect model of COVID-19 pathophysiology, integrating data from over 160 research articles [40]. It contains more than 4,000 nodes and 10,000 relationships representing biological entities such as proteins, genes, chemicals, and biological processes. This graph primarily focuses on host-pathogen interactions, including viral invasion, immune response, comorbidities, and drug-target interactions, making it a vital resource for understanding the virus’s biological impact and therapeutic strategies.

In contrast, the NeuroMMSig KG, first developed by the same group in 2017, focuses on NDDs like AD and PD [41]. This mechanism-enrichment graph is encoded in BEL [42] and represents causal relationships between genes, proteins, and biological processes, integrating multimodal data such as genetic, epigenetic, and imaging features. The graph enables the exploration of disease mechanisms and the identification of potential drug targets by comparing molecular pathways involved in AD and PD.

These COVID-19, AD, and PD KGs, referred to as SCAI-DMaps, are used for further analysis in this study.

#### 2.4.2 Text mining and Natural Language Processing Pipelines

##### 2.4.2.1 Sherpa Kairntech

In this study, we utilized the Sherpa workflow developed by Kairntech [43], for text-based information extraction. Sherpa is a user-friendly, web-based machine learning platform that supports various NLP tasks, including entity recognition and relation extraction using entity fishing (https://github.com/kermitt2/entity-fishing) and openNRE package (https://github.com/thunlp/OpenNRE), respectively. It has been previously employed in our work to extract biological relationships and encode them in BEL [38], [39]. Sherpa was initially trained on a dataset comprising 39,099 distinct triples, constructed by integrating several expert-curated, published KGs covering a range of diseases, including COVID-19, AD, PD, and epilepsy, as described in our previous work [38]. To evaluate Sherpa’s performance on unseen data, we randomly withheld 10% of the training dataset to create an independent test set. The evaluation results are summarized in Table 1 [38]. Sherpa was further updated with a curated dataset on COVID-19 and NDDs compiled by experts at CBM, for use in our current study. The platform achieved an accuracy of 93% after training on this dataset, which included more than 3,000 BEL triples, ensuring high-quality extraction of relationships between biomedical entities for further analysis.

**Table 1.** Performance metrics of the Sherpa text mining workflow for relation extraction on a held-out test dataset. [38].

| **Relation Type** | **Precision** | **Recall** | **F1-score** |
| --- | --- | --- | --- |
| Increase | 74% | 62% | 67% |
| Decrease | 59% | 66% | 62% |
| Directly Increase | 60% | 33% | 42% |
| Directly Decrease | 60% | 42% | 49% |
| Regulate | 66% | 61% | 63% |
| Association | 65% | 54% | 58% |
| Positive Correlation | 64% | 61% | 62% |
| Negative Correlation | 43% | 53% | 47% |
| Biomarker For | 66% | 40% | 50% |
| Has Component | 100% | 33% | 50% |
| Is A | 61% | 30% | 40% |
| Cause No Change | 34% | 37% | 35% |

##### 2.4.2.2 PubTator3

We also leveraged PubTator 3.0 [44], an advanced AI-driven tool developed by the National Library of Medicine (NLM), to extract comprehensive biomedical information from scientific literature. PubTator processes PubMed abstracts and full-text articles, identifying key biological entities and their relationships across various entity types such as chemical–disease, gene-disease, and chemical–gene. Applying PubTator 3.0 API, we extracted annotations in .xml format, which were then parsed and processed into triples (entity-relationship-entity) and saved in an Excel file for further analysis in the study. More complementary information about information extraction with PubTator is added to the Supplementary File, section 5.

### 2.5 Visualization of KGs and Graph Algorithms

To facilitate visualization, navigation, and querying of KGs, we employed an open-source version of the Neo4j platform. Its advanced visualization features enable clear representation and differentiation of various node and relationship types, such as diseases and genes. This capability provides an intuitive and comprehensive view of complex biological interconnections, enhancing both analysis and interpretability. Additionally, Neo4j’s advanced capabilities, such as the Cypher query language and the integration of powerful libraries like the Graph Data Science (GDS) library and the Awesome Procedures on Cypher (APOC) library, facilitate efficient querying and manipulation of graph data. Since the Sherpa tool identifies entity IDs extracted from various databases, such as MeSH, we implemented a post-processing step within the Sherpa text mining pipeline to ensure uniformity and compatibility. This step standardizes and normalizes the extracted entities to align with predefined namespaces and ontologies, ensuring consistency and facilitating interoperability. Details of this post-processing workflow are provided in the Supplementary File, sections 6 and 7. Once processed, all the resulting triples from all KGs were systematically loaded into the Neo4j database for further analysis and exploration.

## 3 Results

### 3.1 General Overview of the Extracted Co-morbidity KGs

The visualization and analysis of KGs from various data sources, as depicted in Figure 3., offers valuable insights into both the structural characteristics of the KGs and the specific biological contexts they aim to represent. Node types were defined based on the subject and object in each triple and were categorized using standard namespaces and ontologies such as HGNC (for genes) [45], Disease Ontology (DO, for diseases) [46], and CHEBI (for chemicals) [47]. Table 2 and Figure 4. provide a comparative overview of several KGs, highlighting differences in the number of nodes, triples, and their densities.

**Figure 3.**
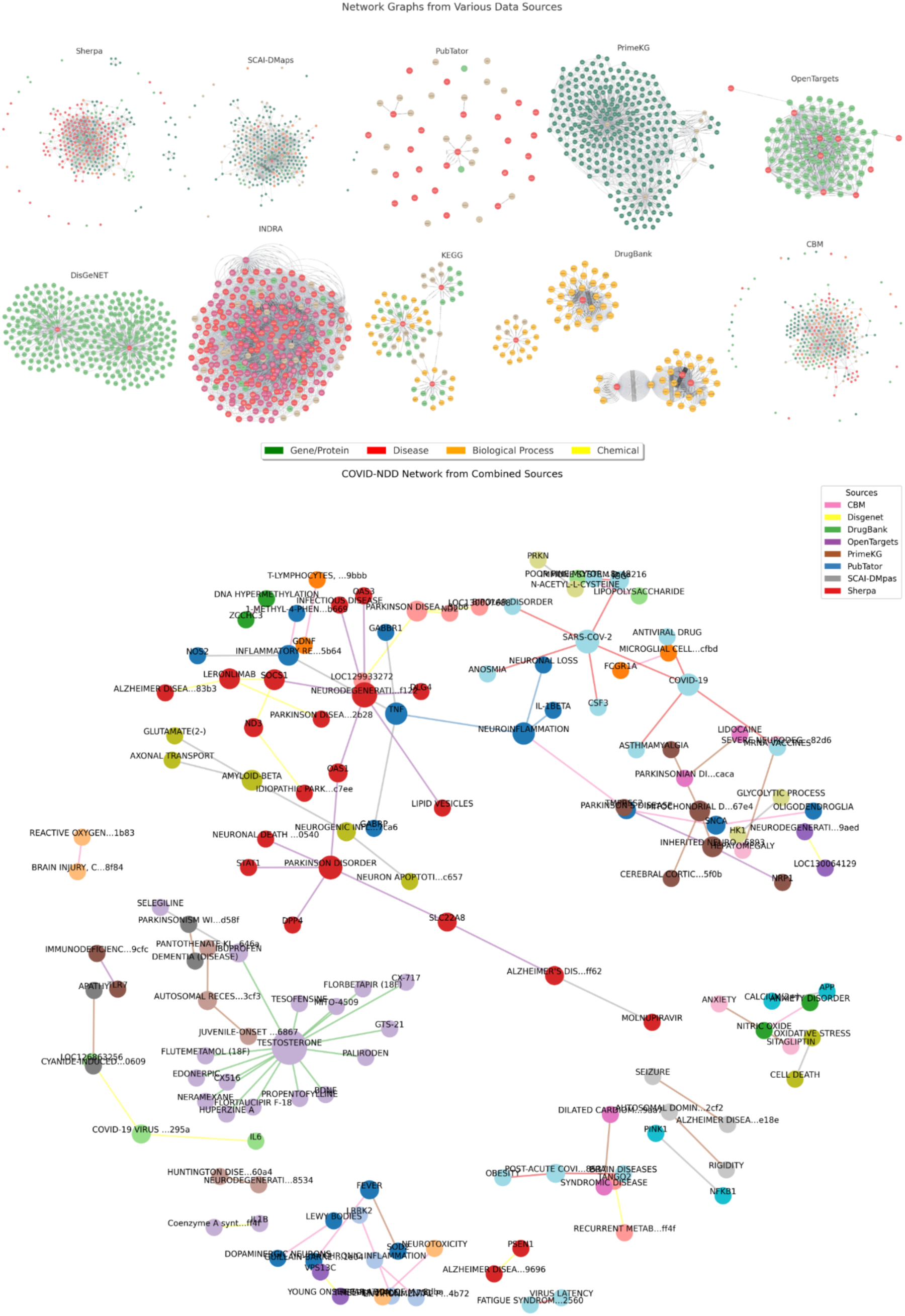
*(Top)* Graph representations in Neo4j illustrating the extracted KGs from individual data sources. Different node types — including genes, proteins, diseases, pathologies, and others — are depicted with distinct colors for improved interpretability. *(Bottom)* Comprehensive community-specific view of various graphs, highlighting how triples are interconnected across various sources. Edge colors correspond to the data source of each triple (as indicated in the legend), while node colors represent the detected clusters and communities within the graph.

**Table 2.**
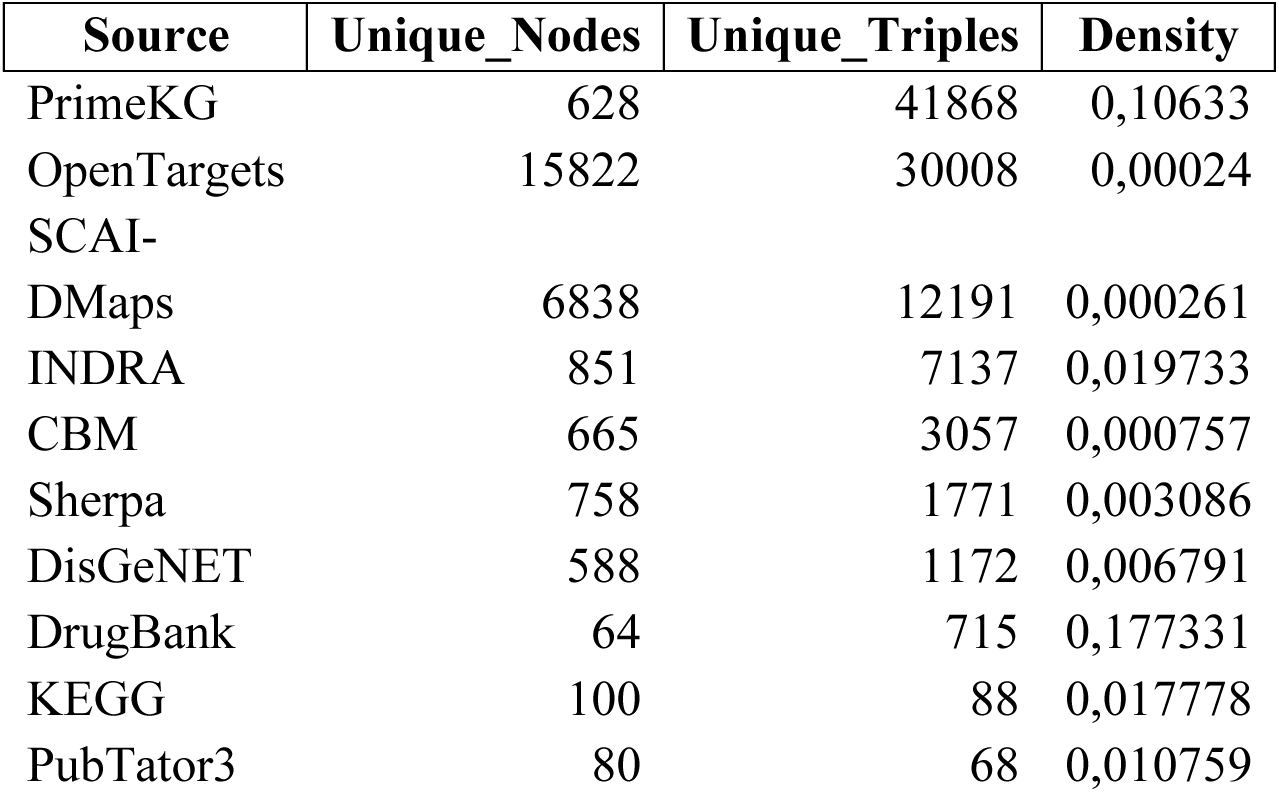
General overview of extracted KGs.

| Source | Unique_Nodes | Unique_Triples | Density |
| --- | --- | --- | --- |
| PrimeKG | 628 | 41868 | 0,10633 |
| OpenTargets | 15822 | 30008 | 0,00024 |
| SCAI-DMaps | 6838 | 12191 | 0,000261 |
| INDRA | 851 | 7137 | 0,019733 |
| CBM | 665 | 3057 | 0,000757 |
| Sherpa | 758 | 1771 | 0,003086 |
| DisGeNET | 588 | 1172 | 0,006791 |
| DrugBank | 64 | 715 | 0,177331 |
| KEGG | 100 | 88 | 0,017778 |
| PubTator3 | 80 | 68 | 0,010759 |

**Figure 4.**
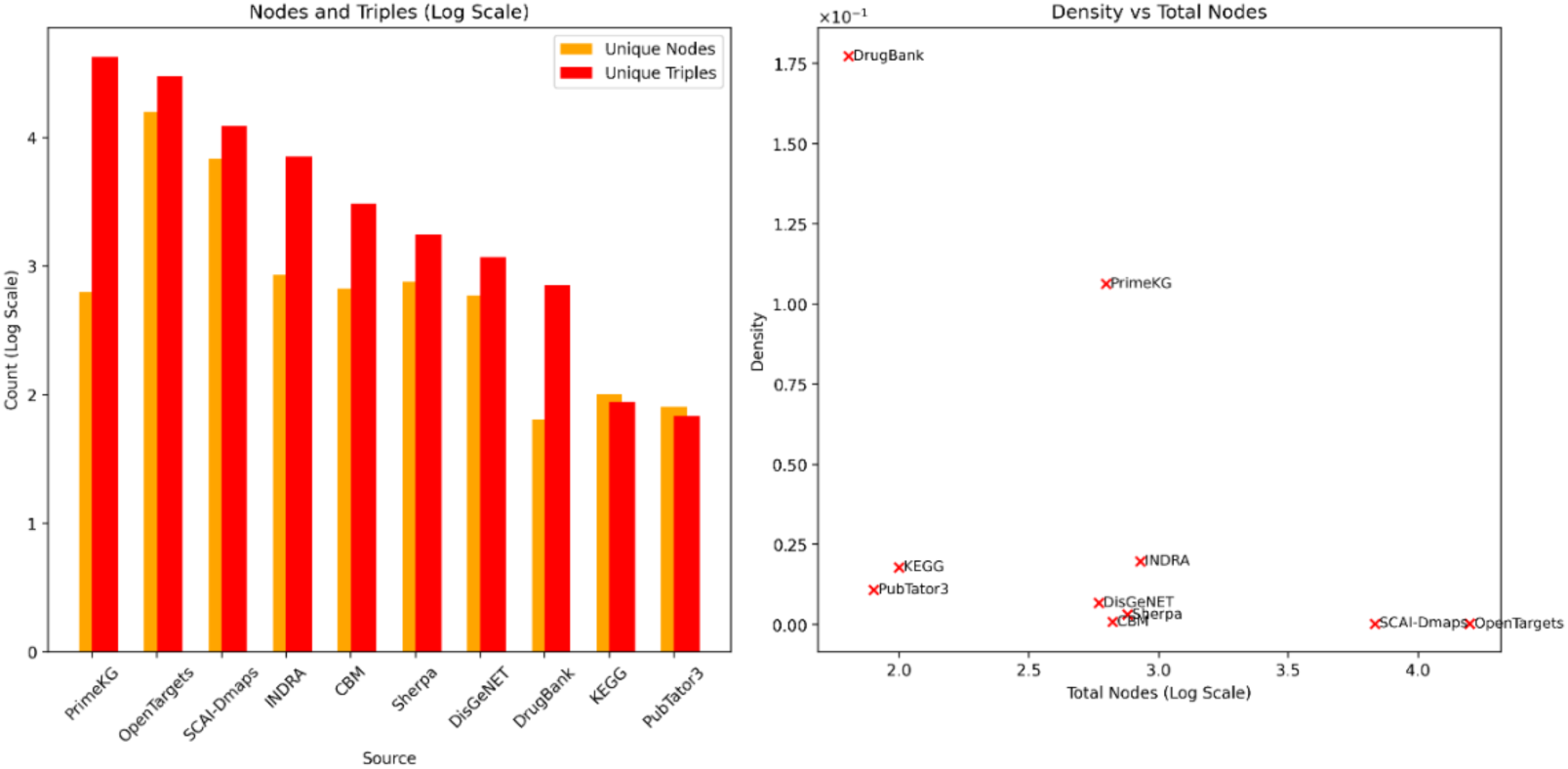
Comparative analysis of COVID-NDD knowledge across different sources. Bar charts (logarithmic scale) show the number of unique nodes and semantic triples from each source, while the scatter plot displays their graph densities relative to total node count, revealing the scale and connectivity patterns of COVID-NDD relationships in each knowledge base.

Focusing on the subgraph of PrimeKG centered on both COVID-19 and NDDs—comprising 628 unique nodes and 41,868 distinct triples—it becomes evident that this versatile graph is designed for scalability and comprehensive analysis. Its relatively high density of 0.1063 reflects a much denser graph structure compared to many others, indicating a tightly connected network of relationships across many entities. Similarly, other large-scale sources like OpenTargets (15,822 nodes, 30,008 triples) and SCAI-DMaps (6,838 nodes, 12,191 triples) present sparse structures, but OpenTargets has a slightly higher density (0.0002397) than SCAI-DMaps (0.0002607). These KGs are designed to cover broad biological domains, offering extensive information with less dense interconnections. Notably, SCAI-DMaps integrates evidence from over 160 peer-reviewed sources, emphasizing host–pathogen interactions and relationships such as increases, associations, decreases, and immune responses. Its construction through manual curation further enhances interpretability and trustworthiness—qualities particularly valuable for exploring inflammation-related comorbidities central to COVID-19.

The CBM graph, with 665 unique nodes and 3,057 triples, provides a moderately sized but sparse KG, with a density of 0.000757. This suggests that while it covers a significant range of entities and relationships, the connections are spread thinly across the dataset. It strikes a balance between scale and specificity, offering a resource that is more expansive than highly specialized graphs like DrugBank, but less dense. CBM is well-suited for exploring broad patterns or integrating with other KGs to generate enriched networks.

In contrast, DrugBank provides a smaller but much denser graph. With only 64 nodes and 715 triples, DrugBank has the highest density (0.1773), indicating a tightly connected graph with a higher number of relationships per node. This focused structure likely represents highly curated information, especially about drug-disease relationships. DisGeNET, another smaller source (588 nodes, 1,172 triples), demonstrates a relatively higher density (0.0067911), highlighting its specialized focus, particularly in gene-disease associations. Both DrugBank and DisGeNET serve as dense, focused KGs that deeply explore disease-specific relationships.

The Sherpa text mining graph, with 758 unique nodes and 1,771 relationships, presents a moderately sized yet specific dataset. Its moderate density of 0.0030864 suggests that it balances meaningful relationships with coverage. Sherpa focuses on regulatory mechanisms and intervention strategies, bridging the gap between the sparsity of large graphs like PrimeKG and the dense connectivity of DrugBank.

When merging different sources into a combined KG, a broader and more comprehensive network emerges. For example, combining data from sources like DisGeNET, OpenTargets, DrugBank, and INDRA results in a graph with a significant number of unique nodes and relationships, especially with OpenTargets contributing 15,822 unique nodes and 30,008 triples. This merged KG offers an expansive view, integrating information about gene-disease associations, pharmacological pathways, and potential interventions, particularly in the context of COVID-19 and neurological complications.

In terms of specificity, the KEGG graph contains fewer unique nodes (100) and triples (88), but is highly curated, focusing on well-established metabolic and signaling pathways. PubTator3 text mining graph, as well, is a small KG derived from biomedical literature, with only 80 unique nodes and 68 relationships, derived mainly from term co-occurrence frequency in scientific research papers related to COVID-19 and NDDs.

The structural diversity across these KGs—ranging from the sparsity of large-scale graphs like PrimeKG and CBM to the density of DrugBank—illustrates the variation of the resources investigated in co-morbidity analysis. Whether offering a broad, exploratory framework (PrimeKG, OpenTargets) or a detailed, specialized network (DrugBank, DisGeNET), each KG provides unique insights into biological interactions in both COVID-19 and NDDs. The figures representing the top 10 nodes with the highest centrality degree for each data source are provided in the Supplementary File. As depicted in these figures, several key nodes reveal important insights into potential comorbidities between neurological disorders and other conditions. “Spinocerebellar ataxia” in PrimeKG emerges as a central node, underscoring its significant role in both motor dysfunction and neurodegenerative comorbidities, suggesting that its involvement might extend beyond movement disorders to other systemic conditions. Additionally, “Fatigue”, a prominent symptom in PrimeKG, appears as a central node, potentially linking a wide range of neurological and systemic diseases, including those associated with viral infections like COVID-19, thus highlighting fatigue as a common underlying feature in co-morbidity networks. In SCAI-DMaps, “APP” and “Amyloid-beta” represent critical molecular mechanisms involved in AD, potentially connecting the pathophysiology of AD with other neurological disorders and systemic conditions, suggesting that these biomarkers may play a role in disease interactions.

Furthermore, the centrality of “Inflammation” and “Neuronal loss” in PubTator3 and Sherpa highlights the crucial role of inflammation as a driver of comorbidities between COVID-19 and NDDs providing insights into how inflammatory pathways could exacerbate disease progression and worsen patient outcomes. Finally, Wolff-Parkinson-White Syndrome, identified in DisGeNET, suggests the possibility of an overlap between cardiovascular and NDDs, indicating the complex nature of comorbidities and their potential interdependence. This insight points to the need for further exploration into how cardiovascular dysfunction might exacerbate neurological conditions, providing a deeper understanding of disease interactions and informing clinical management strategies for patients with multiple comorbidities. DisGeNET exhibited higher node density around well-established risk genes, notably APOE and TREM2, highlighting its genetic disease focus. In contrast, DrugBank formed central hubs around COVID-19 therapeutic agents such as remdesivir and baricitinib, underscoring its pharmacological orientation. Meanwhile, the networks derived from CBM and Sherpa were more abstract and linguistically driven, with dominant nodes labeled ‘BioConcept’, ‘Pathology’, and ‘GeneticFlow’, reflecting their emphasis on text-mined associations and mechanistic interpretations.

### 3.2 Shared Pathophysiology Pathways between COVID-19 and NDD using Shortest Path Analysis

The first step in investigating the interactions between COVID-19 and NDDs was to perform shortest path analysis, aimed at uncovering direct or indirect connections between the two conditions across various graphs. For this means, we developed a custom Cypher query to address the challenges posed by the variations in node names and namespaces across different graphs. This query is designed to uncover connections between COVID-19 and NDDs through a series of structured and methodical steps, as outlined in the Supplementary File, section 8. A central aspect of our analysis was determining the shortest paths between nodes representing various forms of COVID-19 (e.g., COVID-19, SARS-CoV, and COVID) and NDDs (e.g., Alzheimer, Alzheimer’s, Parkinson, Parkinson’s, etc.), while accommodating variations in path lengths or hops. The outcomes of this analysis are summarized in Table 3. Initial findings showed that in the KEGG pathway database, no direct or indirect connections existed between COVID-19 and NDD-related genes, drugs, or pathways. Figure 5. visualizes this path analysis across multiple co-morbidity KGs. The combined dataset of DisGeNET, OpenTargets, DrugBank, and INDRA, demonstrated the highest levels of connectivity and the most diverse pathways linking two diseases. PrimeKG ranked second, still showcasing a strong degree of connectivity. These patterns highlight that different KGs vary in their connectivity pattern and require varying step lengths to establish links between COVID-19 and NDD-related entities. CBM represents a middle ground in terms of connectivity, exhibiting steady, moderate growth across hop levels, indicating a stable network with limited expansion potential. In contrast, graphs from SCAI-DMaps and Sherpa show constrained growth, with minimal increases in unique nodes and paths, making them more suitable for focused, targeted analyses rather than studies requiring extensive connectivity. PubTator3, however, has a highly limited co-morbidity network, showing no expansion beyond its initial connections, underscoring its constrained utility for broader co-morbidity analysis.

**Table 3.** Shortest path analysis between COVID-19 and NDDs nodes using various path lengths (hops).

| Sources | Unique Nodes (Hop = 3) | Unique Paths (Hop = 3) | Total Paths (Hop = 3) | Unique Nodes (Hop = 5) | Unique Paths (Hop = 5) | Total Paths (Hop = 5) |
| --- | --- | --- | --- | --- | --- | --- |
| DisGeNET-OpenTargets-DrugBank-INDRA | 62 | 195 | 208 | 97 | 798 | 898 |
| PrimeKG | 57 | 64 | 64 | 169 | 413 | 415 |
| CBM | 25 | 38 | 39 | 32 | 66 | 67 |
| SCAI-DMaps | 17 | 14 | 14 | 22 | 44 | 44 |
| Sherpa | 3 | 6 | 7 | 6 | 10 | 11 |
| PubTator3 | 1 | 1 | 1 | 1 | 1 | 1 |

**Figure 5.**
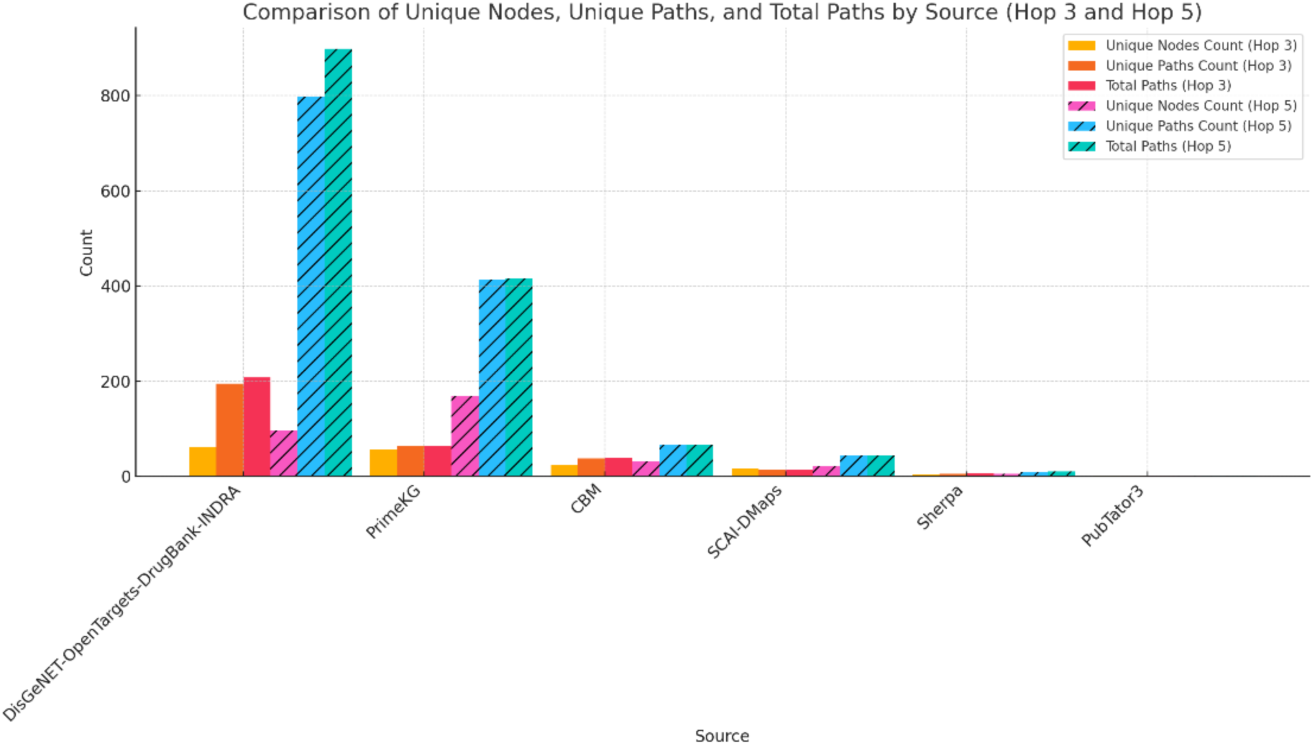
Bar chart summarizing pathway analyses across each co-morbidity KG. For a more comprehensive view, data from the following four sources were integrated for analysis: DisGeNET, OpenTargets, DrugBank, and INDRA. The KEGG pathways database did not reveal any pathways connecting COVID-19-related genes, drugs, or networks.

### 3.3 Exploring Phenotypic and Clinical Endpoint Coverage in Various Graphs Using Human Phenotype Ontology (HPO) Terms

To validate and evaluate the representation of phenotypes within the shortest paths connecting COVID-19 and NDDs, we first identified genes associated with each disease from DisGeNET. Using the Human Phenotype Ontology (HPO) [45], we mapped these genes to their corresponding phenotypic terms, extracting a set of potentially shared phenotypes between COVID-19 and various NDDs. To enhance phenotype matching accuracy within our KG, we cross-referenced HPO terms with synonyms from the MeSH database, enabling a comprehensive and flexible fuzzy matching approach. This method facilitated the identification of nodes in paths that loosely correspond to phenotype terms or their variants, capturing a broader spectrum of relevant phenotypes.

Recognizing limitations in the initial string-based matching approach, we performed additional curation and validation on a benchmark set of 250 path-to-phenotype matches. Each match was reviewed by domain experts to distinguish valid mappings from false positives. This evaluation revealed a low precision of 3.6% (9 valid matches), confirming that string similarity alone can lead to substantial semantic misalignment—for example, incorrectly matching “parkinson” with epilepsy-related phenotypes, which are biologically unrelated. This highlights the need for more sophisticated semantic similarity measures and careful entity selection.

Valid matches consistently exhibited higher similarity scores (mean = 0.672), exemplifying that semantic approaches can succeed when applied to appropriate entity-phenotype pairs. Notably, data source characteristics impacted matching quality: CBM-comorbidity showed the highest precision (33.3%), followed by PrimeKG (3.0%), whereas sources focused on molecular-level associations (OpenTargets, DisGeNet, DrugBank) showed no validated matches. This discrepancy likely reflects differing focuses on phenotypic versus molecular information within these datasets.

Finally, we applied semantic similarity analysis using BERT embeddings to identify and retain only those phenotypes that were likely valid matches, combining automated semantic embedding with manual curation. The network of covered phenotypes is represented in Figure 6. Following this, PrimeKG exhibited the most extensive and validated phenotype coverage, particularly for sensory impairments such as hearing loss and sensory seizures, as well as neurological disorders including childhood-onset neurodegeneration with cerebellar atrophy, hippocampal apoptosis (AD-related), seizures, and sleep disturbances. CBM and Sherpa datasets show narrower but more focused phenotypic profiles, with CBM emphasizing neurological impairments like seizures and motor neuron dysfunction, and Sherpa reflecting more mechanistic molecular phenotypes.

**Figure 6.**
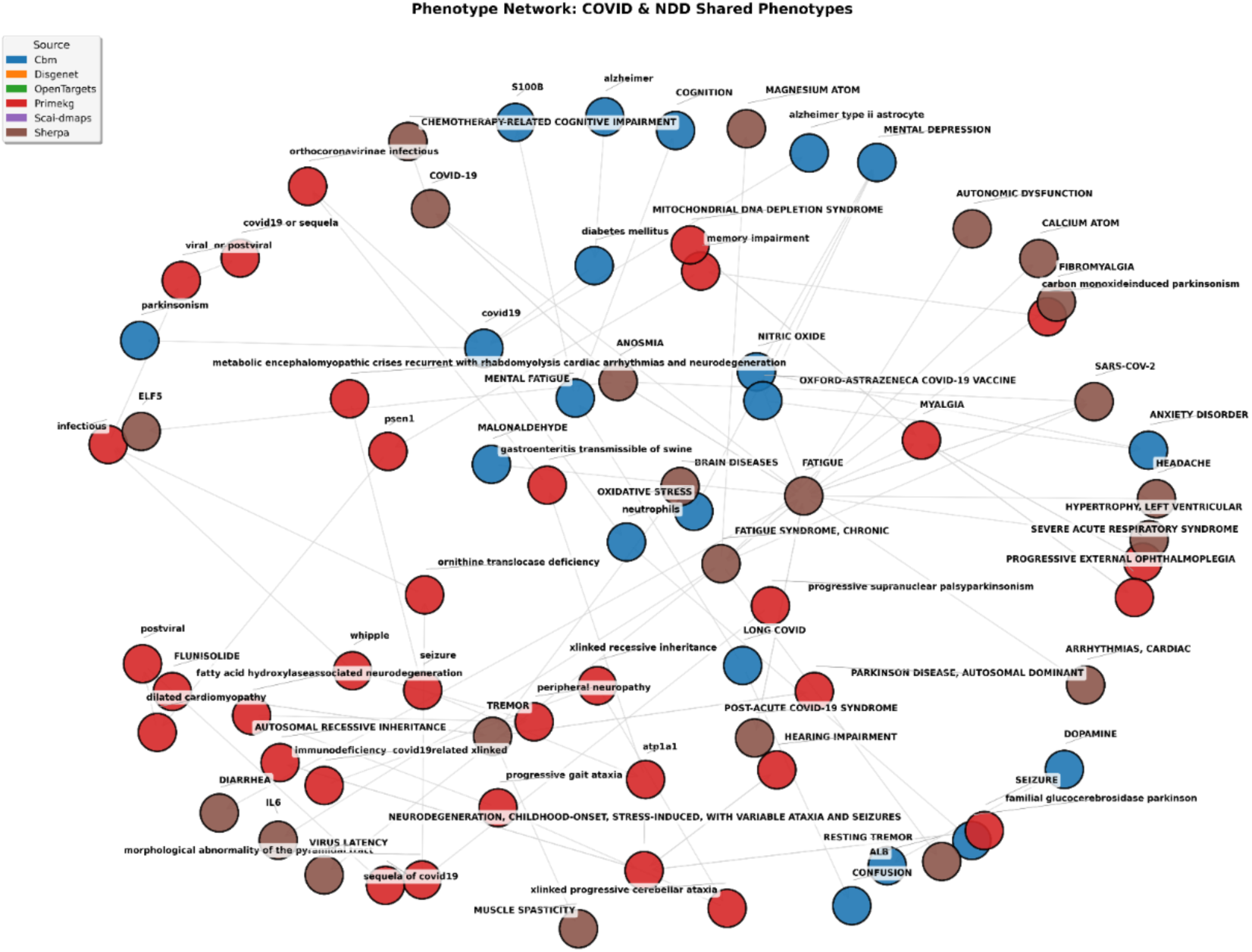
The shared phenotypes between COVID-19 and NDD captured by different sources.

Overall, these results underscore both shared and unique phenotypic landscapes across KGs and emphasize the critical need to integrate diverse, carefully curated datasets and semantic similarity methods to robustly capture the complex phenotype associations linking COVID-19 and NDDs.

### 3.4 Construction of the Co-morbidity Hypothesis Database

#### 3.4.1 Co-morbidity Criteria

To investigate the comorbid relationship between COVID-19 and NDDs, we applied a comprehensive strategy based on several criteria to detect relevant triples and pathways from multiple databases and text mining approaches. The first criterion, direct co-occurrence evidence, was defined as the simultaneous presence of COVID-19 and an NDD within the same path within a maximum of 3 hops. The paths extracted this way were then curated and validated manually and added to the hypothesis space. In the second criterion, shared phenotypic outcomes, we identified clinical symptoms or complications common to both diseases, such as “cognitive decline” or “seizure”, with data sourced from the HPO, as formerly described. In fact, by constructing a curated list of common phenotypes, we aimed to identify biological pathways that reveal functional or mechanistic overlaps at the non-molecular level. This process was complemented by independent literature reviews to collect supporting evidence, including associated genes, protein pathways, cell types, genetic risk factors, and biological processes. To ensure accurate pathway identification, we utilized Neo4j’s Levenshtein fuzzy matching tool, enabling precise recognition of nodes and pathways despite variations in terminology across datasets. The third criterion, common pathophysiological mechanisms, focused on overlapping biological pathways, such as “neuroinflammation”, “oxidative stress”, and “mitochondrial dysfunction”, found in both COVID-19 and NDDs. This was supported by looking through pathway databases like KEGG and Reactome (https://reactome.org), as well as reviewing literature and experimental studies, with examples like IL-6-mediated inflammation and its contribution to both COVID-19 and NDDs [48]. The fourth criterion, shared genetic susceptibility, involved identifying genetic markers linked to both diseases. For instance, the genes NPR3 and TLR7 have been associated with both neurodevelopmental disorders and COVID-19, as evidenced by data from DisGeNET and OpenTargets. The final criterion involved a thorough manual review of potential comorbid mechanisms conducted by domain experts. This step ensured a comprehensive evaluation of the proposed relationships and their biological plausibility. These criteria led to the creation of a co-morbidity database, which integrates these findings into a robust resource for understanding the interaction between COVID-19 and NDDs, offering insights into shared pathophysiological processes, genetic markers, and clinical outcomes. To ensure interoperability, candidate co-morbidity hypotheses were harmonized by standardizing entity names across multiple namespaces and ontologies, such as MeSH, ChEBI, Disease Ontology (DO) (https://disease-ontology.org) and Ontology Lookup Service (OLS) (https://www.ebi.ac.uk/ols4). The harmonized hypotheses were integrated into a publicly available Neo4j Aura database, enabling researchers to explore, curate, and experimentally validate the data. Figure 7. Represents a snapshot of this database.

**Figure 7.**
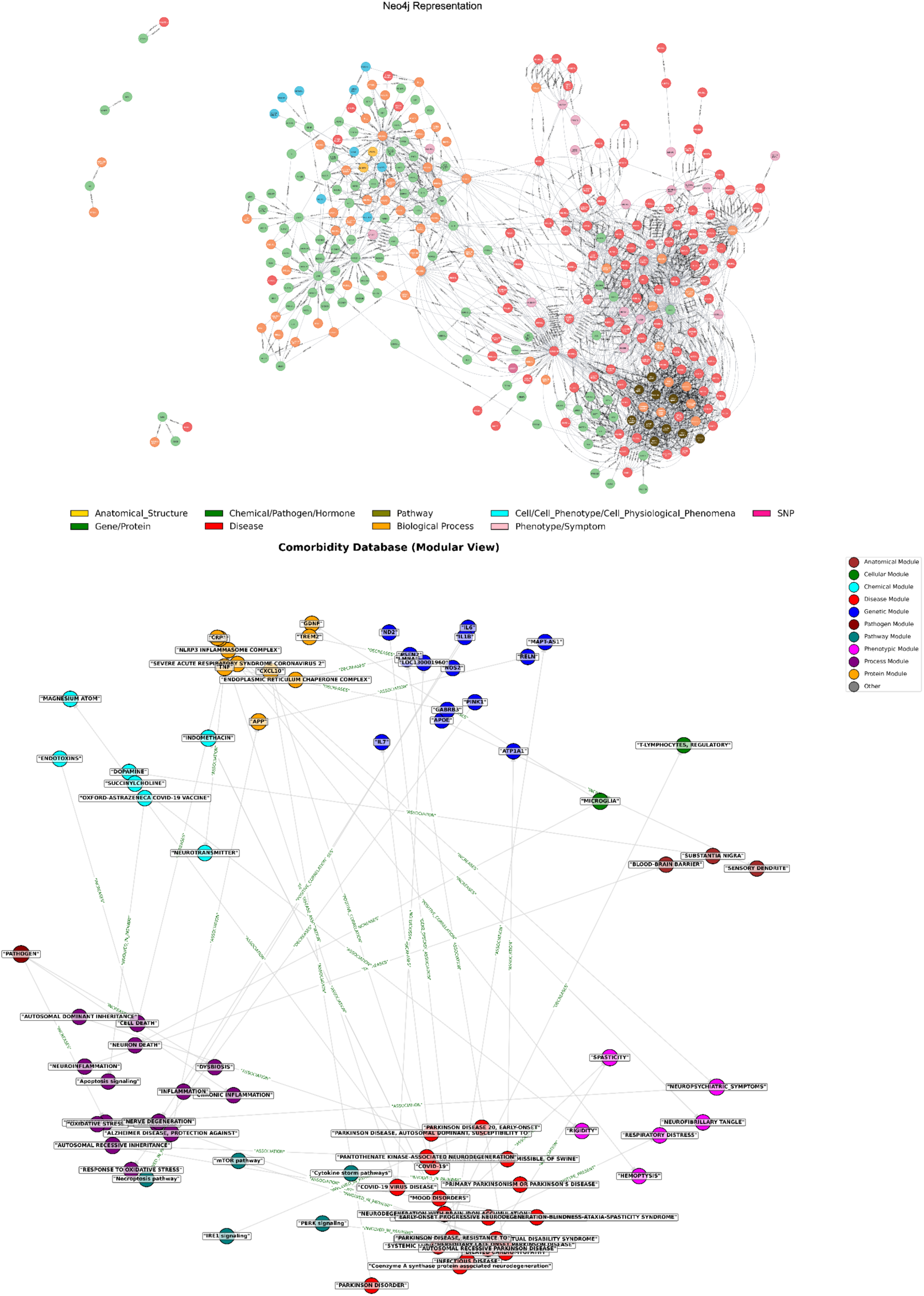
(Top) Overview of the hypothesis database in Neo4j. Various node types are represented by different colors. (Bottom) Integrated modular network displaying the interconnected architecture of genetic, phenotypic, molecular, cellular, anatomical, and other relevant biological modules in the context of COVID-19 and NDD comorbidity.

#### 3.4.2 Enrichment Analysis and Insights of the Co-morbidity Database

To enrich the hypothesis network with genetic data, we integrated risk variants and their documented associations with both COVID-19 and NDDs. This integration was based on data from genome-wide association studies (GWAS), accessible through GWAS Catalog (https://www.ebi.ac.uk/gwas/), which provides a comprehensive database of genetic loci linked to various diseases and traits. Additionally, we supplemented this data with evidence extracted from scientific literature, highlighting established and emerging connections between these genetic risk variants and the two conditions [49], [50], [51]. This approach ensured a more robust and comprehensive framework, enhancing the biological relevance of the hypothesis network by combining high-throughput genetic data with curated research findings. Notably, two specific risk variants, rs5117-C, and rs13107325-T, have been identified as significant single nucleotide polymorphisms (SNPs) associated with PD, AD, and COVID-19, emphasizing shared genetic susceptibilities among the disease, as documented in GWAS studies [52], [53], [54], [55].

The hypothesis network is composed of more than 1,400 nodes and 4,600 edges, with a density of 0.002. The network is predominantly composed of nodes categorized as “GENE,” “DISEASE,” “BIOLOGICAL PROCESS,” “CHEMICAL,” and “PHENOTYPE.” The most frequently observed relationship types within the database include “ASSOCIATION,” “INCREASE,” “INVOLVED_IN_PATHWAY,” “POSITIVE CORRELATION,” and “DECREASE.” The total bar charts illustrating the distribution of relationship types, node types, and namespace types are provided in the Supplementary File.

To assess the relevance of the constructed hypothesis database and investigate the genetic intersections between COVID-19 and NDDs, we conducted a comprehensive manual review of multiple studies [56], [57], [58], [59], [60], [61], [62], [63], [64], [65], [66], [67]. This effort aimed to identify specific genetic variants, polymorphisms, and risk loci that may influence disease susceptibility and progression, offering deeper insights into potential shared biological mechanisms. Our analysis uncovered significant genetic associations between COVID-19 severity and NDDs, all integrated into the constructed hypothesis database. Notable genes associated with increased COVID-19 severity include TMPRSS2 and Furin [60] identified through Sherpa, and TLR7, identified through CBM. Among genes linked to NDDs, Sherpa identified SNCA, whose variants are associated with early-onset PD characterized by dementia and pyramidal signs [66]. Furthermore, PINK1, whose mutations represent a monogenic cause of PD, particularly in early-onset forms [66] was detected via SCAI-DMaps, DisGeNET, and CBM. The AD-associated gene ABCA7, extensively documented in the literature for its SNPs linked to Alzheimer’s, was prominently highlighted by DisGeNET [59]. Several genes were specifically connected to the COVID-19–neurodegeneration link, including NLRP3 [67] as well as OAS1, and CXCL10 [65]. These genes were identified through multiple sources and databases, including CBM, SCAI-DMaps, DisGeNET, and OpenTargets.

Finally, we employed Over-Representation Analysis (ORA) to determine whether known biological pathways are significantly enriched in our gene list [68]. To achieve this, we utilized the DAVID bioinformatics tool (https://davidbioinformatics.nih.gov/), inputting our list of gene symbols, after which DAVID computed the Fisher’s Exact Test and p-value [69], and generated an Enrichment Score. To ensure fair and accurate visualization, we removed statistical outliers and presented the results in a bar chart for clarity.

As illustrated in Figure 8., the pathway enrichment findings reveal key biological mechanisms linking COVID-19 and NDDs. The Nitric Oxide (NO) Signaling Pathway suggests that SARS-CoV-2 may disrupt vascular and neuronal function, contributing to cognitive impairment and neuroinflammation [70], [71]. The NF-κB Activation Pathway, a central regulator of inflammation [72], is over-activated in severe COVID-19 cases, paralleling the chronic neuroinflammatory processes observed in AD and PD [73], [74], [75] . Furthermore, the Chemokine Signaling Pathway highlights how persistent immune activation in COVID-19 could exacerbate neurodegenerative progression by enhancing microglial activation and synaptic dysfunction [76].

**Figure 8.**
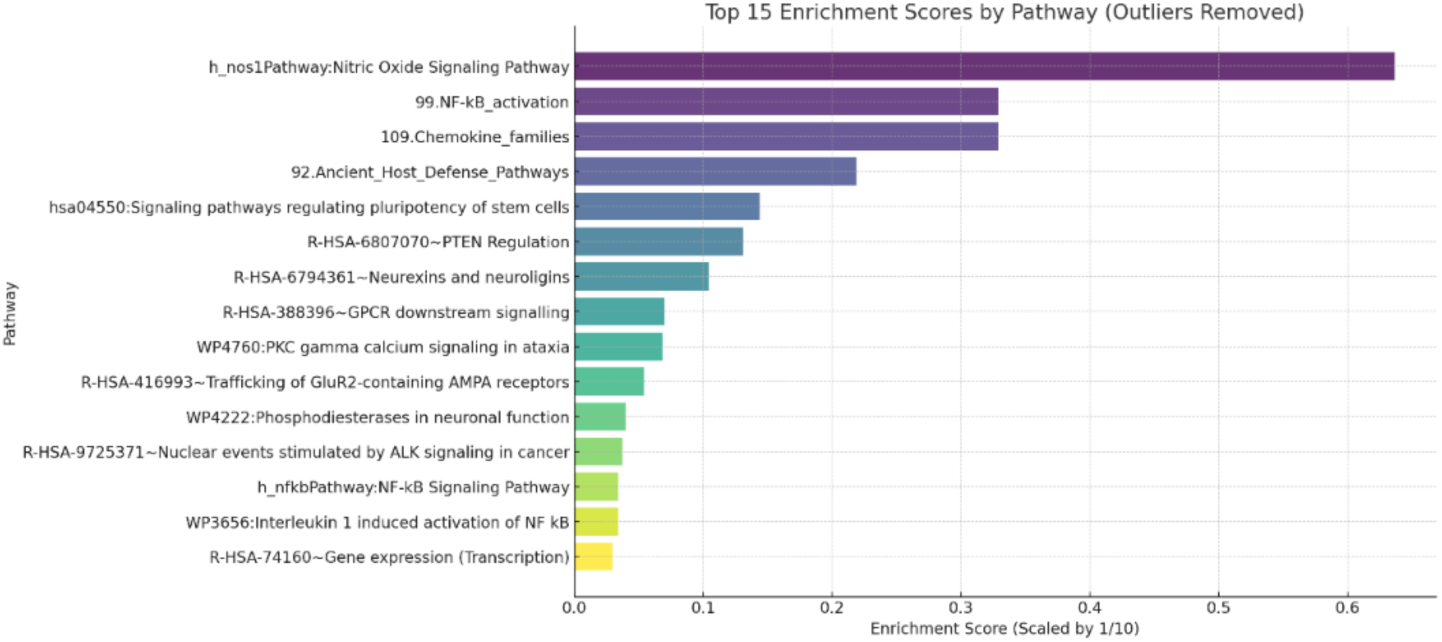
Over-Representation Analysis (ORA) showcasing the top enriched pathways derived from the hypothesis database gene list, analyzed using the DAVID bioinformatics tool.

### 3.5 Validation of COVID–NDD Network Robustness and Connectivity

To ensure the robustness and biological plausibility of the observed connectivity patterns between COVID-19 and NDD entities, we implemented an integrated validation framework combining statistical assessment of path stability with semantic harmonization of terminology. This dual approach was designed to mitigate both methodological artifacts— such as topological biases and random co-occurrences—and lexical inconsistencies arising from heterogeneous biomedical KGs, thereby ensuring the interpretability and reliability of downstream network analyses.

For statistical validation, we designed a comprehensive framework to assess whether the observed network connectivity between COVID-19 and NDD nodes exceeded what would be expected by random chance. We first expanded node identification beyond simple keyword matching by employing semantic similarity analysis using term frequency–inverse document frequency (TF-IDF) vectorization and cosine similarity (threshold = 0.6). This resulted in the identification of 746 COVID-19–related entities and 765 NDD-related entities, selected from curated seed terms encompassing viral mechanisms, neurodegenerative processes, and disease phenotypes.

We conducted five complementary statistical tests across 1,000 permutation iterations to evaluate the connectivity between COVID and NDD nodes. These included assessments of direct connectivity (measuring the number of direct edges between node sets), triple-level co-occurrence (analyzing the co-appearance of COVID and NDD terms within individual KG statements), shortest path analysis (evaluating the distribution of path lengths and counts between node sets), network proximity (measuring average distances between node groups), and degree-preserving connectivity (using edge-swapping algorithms to control for inherent network topology). For each test, we generated empirical null distributions by randomizing node sets while preserving the original network’s degree distribution and node count.

The analysis revealed highly significant connectivity between COVID and NDD nodes, as demonstrated in Figure 9. Direct connectivity showed strong enrichment, with 1,346 observed edges versus 727 expected (p = 0.001; Bonferroni-corrected p = 0.006; Cohen’s d = 6.92), indicating a large effect size. Similarly, the shortest path count test revealed a fivefold increase in observed paths (240 observed vs. 45 expected; p = 0.001; Bonferroni p = 0.006; Cohen’s d = 15.61), demonstrating strong statistical signal. Triple-level co-occurrence analysis yielded 2,921 observed co-mentions—double the null expectation of 1,437 (p = 0.001; Bonferroni p = 0.006; Cohen’s d = 9.94). These three tests remained statistically significant even after applying stringent Bonferroni and FDR corrections (α = 0.05), underscoring the robustness of the observed associations.

**Figure 9.**
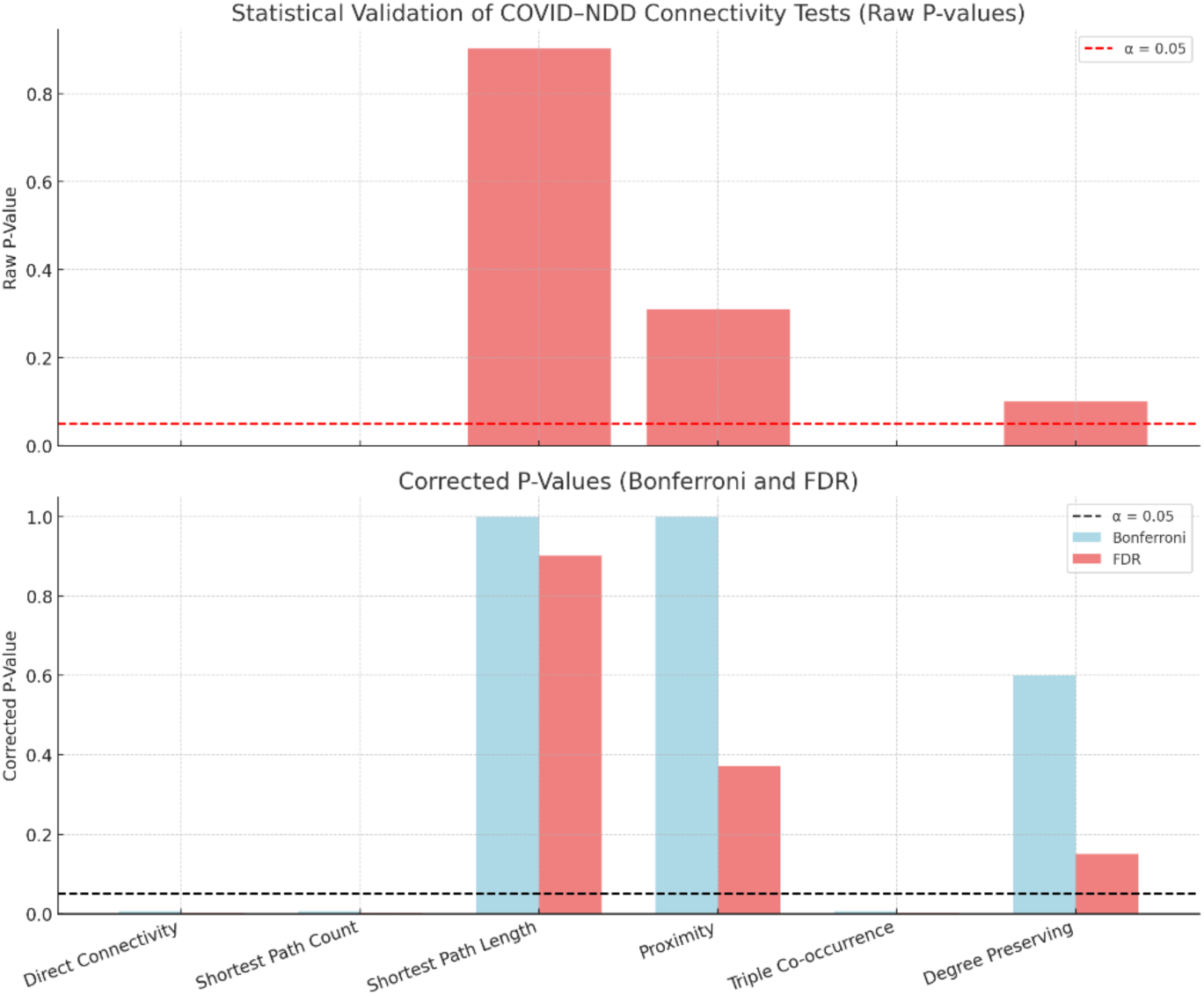
Statistical Validation of COVID–NDD Connectivity Patterns Using Permutation Testing and Multiple Correction Methods. The top panel displays raw p-values for five complementary statistical tests assessing COVID–NDD network connectivity, with the red dashed line indicating the α = 0.05 significance threshold. The bottom panel shows Bonferroni- and FDR-corrected p-values for the same tests. Direct connectivity, shortest path count, and triple co-occurrence tests remain statistically significant after correction, demonstrating robust enrichment beyond chance.

In contrast, topological metrics such as average shortest path length (observed = 4.296 vs. 4.493 expected; p = 0.9021) and proximity-based distance (observed = 4.019 vs. 4.326 expected; p = 0.3107) did not reach statistical significance, indicating that the COVID–NDD connectivity signal is not primarily driven by global graph compactness or shorter indirect paths. Similarly, degree-preserving null model analysis yielded only a modest enrichment of direct COVID–NDD links (1,346 observed vs. 1,318 expected; p = 0.0999), failing to reach correction thresholds. Importantly, these non-significant results reinforce that the strongest comorbidity signal emerges not from generic network architecture, but from biologically grounded relationships—including direct edges, shared molecular entities, and high-confidence co-occurrences—captured through curated data and semantic enrichment.

In parallel, to mitigate the impact of lexical inconsistencies across integrated KGs, we performed a systematic graph harmonization analysis. Disease and phenotype terms were mapped to standardized MONDO identifiers using the EBI Ontology Lookup Service API, following preprocessing steps that included character normalization and removal of punctuation. Lexical variants were identified by computing cosine similarity on character-level TF-IDF vectors, applying a similarity threshold of 0.90 to detect and cluster synonym groups. Each synonym cluster was collapsed to a canonical label, typically the longest variant. Additionally, a curated lexicon of 54 canonical biomedical terms was used to guide fuzzy string matching for enhanced synonym mapping.

The harmonization process identified approximately 8% redundancy across 393 analyzed nodes and consolidated six distinct synonym groups, largely composed of abbreviation variants and spelling alternatives. MONDO integration successfully mapped 27 disease terms to standardized identifiers, including central entities such as Alzheimer’s disease (MONDO:0004975), COVID-19 (MONDO:0100096), and Parkinson’s disease (MONDO:0005180). To quantify the impact of harmonization, we computed Spearman and Pearson correlation coefficients on node frequency distributions before and after harmonization. These analyses demonstrated high rank stability, with average Spearman correlation of 0.998 (range: 0.995–1.000) across KGs, indicating that centrality measures and hub node hierarchies remained intact.

We further assessed the biological stability of pathway-level insights post-harmonization (Supplementary File, section 8). Rankings of 15 key biological pathways and concepts related to COVID-19 and NDDs—such as neuroinflammation, immune response, ACE2, APOE, IL-6, TNF-α, and tau protein—remained within 5% of their original centrality values. This confirms that harmonization did not distort critical mechanistic insights. Notably, path-based analyses performed on harmonized graphs showed consistent results with the original graphs: COVID and NDD nodes remained significantly more interconnected via shortest paths than expected by chance (p < 0.05), and their average path lengths were shorter compared to randomized networks (p < 0.05). These patterns persisted even after controlling for hub-node bias using degree-preserving null models, reinforcing the conclusion that the observed associations reflect biologically meaningful, indirect pathways rather than structural or lexical artifacts.

Together, the integration of statistical validation and harmonization analysis confirms that the identified COVID–NDD comorbidity network captures genuine biological relationships. The robustness of node centrality, pathway preservation, and statistical enrichment across multiple validation schemes strengthens the utility of the constructed KGs for downstream translational research.

### 3.6 Comparison and Alignment with Other COVID-19 Knowledge Sources

To comprehensively situate our work within the current landscape, we compared our approach to three notable computational frameworks in the COVID-19 domain, highlighting several key differentiators:

1. KG-COVID-19 (Reese et al., 2021) [77] : A flexible framework for producing customized KG for COVID-19 response that integrates 13 knowledge sources containing 377,482 nodes and over 21 million edges through an automated three-step process (download, transform, merge). This framework primarily focuses on general drug repurposing and machine learning applications.
2. CIDO (Coronavirus Infectious Disease Ontology) (He et al., 2020)) [78]: A community-based ontology containing over 4,000 terms that provides standardized vocabulary for coronavirus disease knowledge covering etiology, transmission, epidemiology, pathogenesis, diagnosis, prevention, and treatment. CIDO follows OBO Foundry principles and excels at knowledge standardization and semantic interoperability.
3. Network-based drug repurposing framework for COVID-19 treatment targeting central nervous system disorders (Qian et al. (2024) [79]). Their study utilized an integrative computational approach combining comorbidity network construction, modular analysis, and dynamic perturbation methods to identify therapeutic targets and drug candidates specifically for COVID-19 patients with CNS complications. Focusing on four neurological conditions—AD, PD, multiple sclerosis, and autism spectrum disorder—they constructed protein-protein interaction networks from shared genes between COVID-19 and these CNS disorders, applied connectivity map (CMap) drug perturbation analysis to identify target modules, and employed molecular dynamics simulations to validate specific drug-target interactions.

The comparison clearly demonstrates that while these existing frameworks provide valuable foundations for COVID-19 research, our work addresses a specific gap that neither systematically explores: the mechanistic understanding of COVID-19 and NDD comorbidities. Unlike the broad-application scope of KG-COVID-19 and CIDO frameworks, we specifically target COVID-19/NDD comorbidity and interactions—a critical clinical concern given observed neurological complications and increased vulnerability of NDD patients. We developed comorbidity-specific analytical methods (shortest path analysis, phenotype coverage assessment, hypothesis generation criteria) that provide actionable insights for patient care, extending beyond general knowledge organization.

While our study provides comprehensive mechanistic hypotheses through multi-modal data integration and statistical validation, Qian et al. demonstrate a pathway for translating these insights into concrete therapeutic interventions. Specifically, they show that modulating UCHL1 via dopamine-related compounds — notably, using pregnenolone as a dopamine release enhancer — may confer clinical benefit. This finding aligns with our identification of neurotransmitter signaling pathways within COVID-19–NDD networks. Moreover, their focus on elderly patients with CNS disorders as a priority population for these interventions is consistent with our findings regarding age-related susceptibility factors, though our broader framework suggests that demographic stratification should be systematically integrated into comorbidity modeling

Their application of molecular dynamics simulations to validate drug candidates pregnenolone and BRD-K87426499 as UCHL1 modulators, exemplifies how the broad hypothesis space generated by our KG approach can be systematically narrowed to actionable therapeutic targets. The convergence of their focused drug repurposing analysis with our systems-level pathway identification suggests that the most promising therapeutic strategies may emerge from combining comprehensive mechanistic mapping—as provided by our framework—with targeted experimental validation and drug development approaches. This integration addresses a critical gap in computational biomedical research, where broad hypothesis generation often lacks clear pathways to clinical application.

This comparative analysis was particularly valuable because their focused, experimentally-validated approach provides an opportunity to assess whether our broader, multi-graph methodology identifies similar biological mechanisms and therapeutic targets, thereby evaluating the clinical relevance and translational potential of our comprehensive computational framework.

The alignment between our findings and those of Qian et al. establishes a validated framework for investigating complex disease comorbidities that combines the breadth of KG integration with the specificity of network-based drug discovery. Our identification of inflammatory cascades, particularly IL-6/NF-κB activation pathways, provides the broader mechanistic context within which specific targets like UCHL1 operate, while their experimental validation demonstrates that computationally-predicted targets can be successfully translated to candidate therapeutics. This complementary approach addresses limitations inherent in both methodologies: our comprehensive but potentially noisy multi-source integration benefits from their focused experimental validation, while their targeted analysis could potentially gain broader mechanistic context from our systems-level investigation. Future comorbidity research can potentially leverage this integrated approach, using comprehensive KG analysis to generate mechanistic hypotheses followed by focused network-based drug repurposing to identify and validate specific therapeutic interventions.

We propose that future research into disease comorbidities—especially those with complex, age-related etiologies such as COVID-19 and NDDs—will be best served by embracing a hybrid paradigm. Computational frameworks one presented in this study, are well-suited to systematically map the mechanistic landscape, uncovering biologically meaningful associations and prioritizing candidate pathways and molecular targets. These hypotheses can then be experimentally validated and iteratively refined, facilitating both the forward translation of discoveries into therapeutic strategies and the backward optimization of computational models for greater biological fidelity.

Accordingly, our contribution extends beyond the development of a KG database—it establishes a scalable, integrative framework for discovery, in which high-throughput, AI-driven hypothesis generation operates in synergy with targeted, experimentally grounded validation. This approach is particularly powerful in domains characterized by biological complexity, fragmented data ecosystems, and urgent clinical need, offering a path toward more actionable, mechanistically grounded insights.

## 4 Discussion

This study explored the comorbidity between COVID-19 and NDDs by analyzing their relationships within diverse KGs. Using insights from biomedical databases and text mining tools applied to relevant publications, we examined mechanisms and pathways linking COVID-19 to long-term neurological complications. Our approach, which combined graph algorithms with analyses of comorbid mechanisms, endpoints, and symptoms, allowed for a detailed exploration of how COVID-19-related processes intersect with NDD pathways. These findings contribute to the growing body of knowledge on the broader health implications of the virus, particularly its potential to exacerbate or initiate neurodegenerative processes.

The findings from the multi-scale graph analysis provide mechanistically grounded hypotheses regarding the co-morbidity between COVID-19 and NDDs. By identifying shared nodes, paths, and phenotypic features across curated and text-mined KGs, our study surfaces potential biological mediators—such as inflammatory markers, host-pathogen interaction nodes, and neurological phenotypes—that may underlie disease intersection. These insights are actionable: researchers can test specific path hypotheses in vitro, investigate gene expression profiles in patient cohorts, or design targeted omics studies to validate molecular overlaps. Moreover, the shortest-path analyses and centrality rankings highlight candidate genes and pathways for follow-up in mechanistic and interventional studies.

### 4.1 Insights from various Co-Morbidity KGs

The structural diversity of the analyzed KGs, reflected in their distinct densities, connectivity patterns, and thematic focuses, highlights their complementary roles in biomedical research. DrugBank, for example, focuses on high-resolution drug-disease relationships, emphasizing molecular interactions relevant to precision pharmacological targeting. This specificity makes it particularly suitable for identifying therapeutic interventions. On the other hand, broader resources such as PrimeKG and OpenTargets take a systems-level approach, capturing the interdependencies between genes, proteins, and biological processes. This perspective is particularly useful for studying diseases with overlapping mechanisms or shared phenotypic manifestations, such as the complex interplay between COVID-19 and NDDs.

Intermediate-sized KGs offer valuable, targeted molecular insights. SCAI-DMaps uncovers relationships involving neurotrophic factors, such as how GDNF and BDNF influence PD progression (with decreases observed in Parkinson’s). It also highlights immune system responses, revealing the upregulation of IL2 in the context of COVID-19. Sherpa maps the associations between COVID-19 and mood disorders, emphasizing the psychological impact of the virus KGs extracted through CBM and SCAI-DMaps contribute critical mechanistic insights, such as the role of mitochondrial metabolism in PD (as detailed in the study by Kannarkat et al. [80]), the processes governing blood-brain barrier development, and the dynamics of neuroinflammation and cytokine feedback loops.

DisGeNET and UNIPROT contribute to genetic associations (ND4—ASSOCIATION--Alzheimer Disease, mitochondrial), while PrimeKG captures complex phenotypes (X-linked inheritance patterns). Recent COVID-specific insights from CBM graph also highlight Long COVID’s effects on cognition and mental health.

These complementary KG strengths, from molecular mechanisms to systemic interactions, provide a comprehensive understanding of COVID-19-NDD relationships.

### 4.2 Key Genetic and Inflammatory Mediators Driving COVID–NDD Comorbidity

Across analyses, several genetic and inflammatory markers consistently emerged as pivotal in understanding the intersection of COVID-19 and NDDs. Markers such as TLR7 (Toll-like Receptor 7), VEGFA (Vascular Endothelial Growth Factor A), APOE (Apolipoprotein E), and a variety of cytokines highlight the central roles of genetic susceptibility and immune responses. These markers are well-documented as contributors to inflammatory processes that can exacerbate both viral infections and neurodegenerative conditions [81], [82], [83].

APP’s interaction with oxidative stress response pathways provides another mechanistic link, documented in SCAI-DMaps [84]. CBM data emphasizes neuroinflammation’s role in modulating adaptive immune responses, while literature evidence confirms feedback loops between inflammatory cytokine production and neuroinflammation.

Neutrophil activation which is a critical mechanism involved in severe COVID-19 pathophysiology [85], was successfully captured by CBM KG. During infection, activated neutrophils release inflammatory mediators, form extracellular traps (NETs), and produce reactive oxygen species, leading to tissue damage and organ dysfunction [85]. This mechanism, suggests a potential link between acute COVID-19 severity and long-term neurological complications, highlighting its importance as a therapeutic target. Neutrophil activation which is a critical mechanism involved in severe COVID-19 pathophysiology [85], was successfully captured by CBM KG. During infection, activated neutrophils release inflammatory mediators, form extracellular traps (NETs), and produce reactive oxygen species, leading to tissue damage and organ dysfunction [85]. This mechanism, suggests a potential link between acute COVID-19 severity and long-term neurological complications, highlighting its importance as a therapeutic target.

Sherpa further identified TRPV4, a calcium-permeable ion channel, as a notable marker involved in COVID-19 pathophysiology. TRPV4’s association with inflammation, respiratory distress, and cellular stress underscores its potential role as a mechanistic link between acute COVID-19 symptoms and long-term neurological complications [86], [87]. The consistent identification of these markers across KGs reinforces their relevance for further research and highlights their potential as therapeutic targets.

### 4.3 Cross-Disease GWAS Enrichment and Genetic Overlap with Neurodegeneration

To further evaluate the biological relevance of the genes prioritized in our multi-source co-morbidity KGs, we conducted a comprehensive review of publicly available genome-wide association studies (GWAS), with a focus on those curated in the GWAS Catalog. The aim was to identify genetic loci implicated in both COVID-19 susceptibility or severity and neurodegenerative or neuroimmune conditions, thereby reinforcing the genetic basis of comorbidity.

Several genes emerged from this integrative analysis as having dual relevance. IFNAR2, a key interferon signaling gene, appeared consistently in GWAS studies associated with severe COVID-19 outcomes. Although not traditionally linked to neurodegenerative disorders, recent research suggests that dysregulation of interferon signaling may play a role in microglial activation and synaptic degradation, both of which are critical features of AD pathology [88]. Similarly, TNF, a gene encoding tumor necrosis factor—a pro-inflammatory cytokine—was identified in GWAS data for both COVID-19 and Type 1 diabetes. TNF has long been implicated in chronic neuroinflammation and glial cell activation, central mechanisms in both Alzheimer’s and Parkinson’s diseases.

Other interleukins, notably IL2 and IL21, were also found in GWAS datasets related to autoimmune traits and COVID-19 severity. IL2 appeared prominently in our SCAI-DMaps and Sherpa-extracted KGs as a central node within inflammation-related modules. Its known role in T-cell activation and regulation of central nervous system inflammation positions it as a key player in diseases such as multiple sclerosis and PD. MAPT, the gene encoding the tau protein, was found to be directly associated with both PD and AD through its involvement in tauopathy. It also appeared in COVID-19–related GWAS studies and was enriched as a highly connected node in PrimeKG, DisGeNET, and SCAI-DMaps.

The convergence of these GWAS-identified loci with genes prioritized in our comorbidity KGs provides strong independent validation of the mechanistic overlap between COVID-19 and neurodegenerative disorders. This genetic cross-validation reinforces the biological plausibility of shared neuroimmune and inflammatory pathways driving both conditions. As a result, we have integrated these GWAS-supported associations into our comorbidity hypothesis database, further grounding our predictions in clinically relevant genetic evidence and enhancing the translational potential of our findings.

### 4.4 Optimization of Co-Morbidity Modeling

A central objective of this study was to optimize comorbidity modeling in a manner that maximizes both the breadth and diversity of biologically plausible hypotheses—what we refer to in this context as *recall*. Rather than limiting our analysis to a small set of well-characterized associations, our framework prioritizes the comprehensive identification of meaningful relationships among biomedical entities, including genes, diseases, proteins, SNPs, and phenotypes. This expansive approach reflects the exploratory nature of comorbidity hypothesis collection, where the value lies not only in confirming known associations but also in uncovering mechanistic links that can guide hypothesis generation and downstream investigation.

To this end, we implemented a multi-metric evaluation framework to assess the structural, semantic, and functional properties of the constructed and integrated KGs. These metrics provide quantitative evidence for the optimization of graph topology, connectivity, and semantic richness, demonstrating how integration enhances biological relevance and hypothesis discovery capacity.

Graph topology optimization was observed in the balance achieved between compactness and coverage. The integrated KG combined the structural density of drug–disease relationships with broader, mechanistically diverse information from heterogeneous sources. This enabled the graph to retain high information content while remaining navigable. Centrality analyses revealed that key hubs involved in COVID-19–NDD interactions consistently aligned with biologically meaningful entities, further supporting the plausibility and utility of the identified pathways.

Connectivity optimization was reflected in the increased richness and diversity of mechanistic paths between COVID-19 and NDD nodes in the integrated graph, as compared to individual source KGs. This improvement in path diversity translates into enhanced pathway discovery potential and higher recall of functionally relevant connections. By expanding the space of biologically interpretable paths, the model facilitates exploratory traversal through multiple layers of disease biology.

Phenotype coverage optimization was another major advantage of the integrated approach. The final hypothesis database captures a wide spectrum of clinical phenotypes shared between COVID-19 and NDDs, ranging from cognitive decline to systemic inflammatory responses. This more complete phenotypic representation bridges the gap between molecular mechanisms and patient-level manifestations, increasing the translational relevance of the generated hypotheses.

Semantic and mechanistic diversity was significantly enhanced by the inclusion of a broad array of entity and relation types, spanning genes, pathways, biological processes, and clinical outcomes. The integration of heterogeneous data sources enabled the model to capture distinct layers of disease biology—from genetic predispositions and molecular cascades to phenotypic expressions and clinical complications—thereby enabling multi-level hypothesis generation that few individual KGs could support on their own.

Finally, data harmonization into a standardized, ontology-driven schema ensured more semantic consistency and structural interoperability across the integrated KG. This design facilitated seamless querying and enabled complex, multi-hop reasoning over entities and relationships. As a result, the comorbidity graph supports in-depth, cross-scale hypothesis generation traversing genes, pathways, phenotypes, and therapeutic targets—an essential characteristic of optimized recall and modeling efficacy.

Taken together, the unification of complementary resources, optimization of semantic and structural connectivity, and expansion of mechanistic hypothesis space resulted in a robust framework for comorbidity modeling. By maximizing the diversity and plausibility of the associations captured, our approach lays a strong foundation for both exploratory biomedical research and can further aid in the identification of therapeutic opportunities in high-risk patient populations.

### 4.5 Utility of the Comorbidity Database and Drug Repurposing Use Cases

The comorbidity hypothesis database and KG developed in this study provide practical utility for a wide range of stakeholders, bridging molecular insights with clinical and translational applications. The publicly accessible database offers a structured, graph-based framework for hypothesis generation and exploration, enabling researchers to navigate mechanistic and phenotypic pathways that underlie the observed associations between COVID-19 and NDDs. Clinicians can leverage these insights to better understand why individuals with pre-existing NDDs appear at increased risk for severe COVID-19 outcomes, or conversely, how SARS-CoV-2 infection might contribute to the initiation or progression of neurodegenerative processes. This knowledge may inform risk stratification, individualized monitoring strategies, and clinical decision-making for vulnerable patient populations.

Our KG systematically integrates multi-scale biological data and literature-derived hypotheses, and already includes selected drug–target annotations where available in the integrated sources. For example, we identify the association of glucagon-like peptide-1 receptor (GLP-1R) agonists with APOE, a gene strongly implicated in neurodegeneration. This highlights a potentially druggable node in the pathway linking COVID-19-related mechanisms to neurodegenerative disorders, illustrating the value of our graph in pinpointing actionable intervention points. Practitioners can already begin to use the current hypothesis graph as a resource for hypothesis generation and prioritization, extracting mechanistic pathways connecting COVID-19 to neurodegenerative processes, and identifying points of intervention annotated with drug classes such as GLP-1R agonists in APOE-related pathways (Supplementary file, section 10.). This graph-driven approach enables early-stage exploration and refinement of drug repurposing strategies by reducing the search space and increasing the likelihood of identifying translationally relevant targets.

However, it is important to emphasize that the current KG does not yet provide exhaustive drug–target coverage. While some annotations are embedded, many important inflammatory mediators—such as IL-6 and TNF—lack a complete set of known inhibitors within the graph. Future enrichment efforts will involve systematically incorporating comprehensive pharmacological annotations from external drug databases, thereby expanding the utility of the graph for therapeutic prioritization. By overlaying drug–target relationships on mechanistically relevant subgraphs—such as those involving neuroinflammation, mitochondrial dysfunction, or blood–brain barrier disruption—researchers will be able to more thoroughly identify candidate compounds for repurposing and direct experimental validation.

Crucially, validating the mechanistic hypotheses and candidate targets derived from the KG will require access to real-world, patient-level datasets, such as electronic health records, prescription histories, and longitudinal cohort outcomes, in addition to targeted experimental models. Given the scale and complexity of the hypotheses represented in the KG, it is neither feasible nor efficient to experimentally test all candidates through traditional wet-lab pipelines. As such, we advocate for the public release of our annotated hypothesis database to catalyze collaborative validation within the broader research community. This open-access strategy aligns with the mission of the COMMUTE project, which serves as a collaborative framework for harvesting, sharing, and refining comorbidity hypotheses through observational studies, in silico analyses, and iterative data-driven exploration.

### 4.6 Integrating Heterogeneous KGs: Semantic, Structural, and Validation Challenges

A key challenge encountered in this study was the heterogeneity of data across KGs. Each KG is derived from different sources and reflects unique strengths and limitations. For instance, some KGs focus on molecular interactions with high specificity but may lack phenotypic or clinical associations, while others provide broader, systems-level insights that might overlook molecular details. This diversity necessitates the integration of multiple KGs to create a more complete representation of the mechanisms underlying COVID-19 and NDD comorbidities.

Node harmonization is a foundational yet inherently complex step in integrating data from heterogeneous KGs. Standardizing entities by aligning them to reference ontologies ensures semantic consistency and enables meaningful cross-KG comparisons. However, discrepancies in terminologies, identifiers, and classification schemes across datasets pose significant challenges. A single biological concept may be represented under different labels or identifiers, leading to redundancy, ambiguity, or misalignment in downstream analyses. In this regard, to improve the semantic consistency of our network analyses, we implemented an automatic term harmonization step that grouped and collapsed synonymous node labels across all KGs. Using a TF-IDF-based string similarity approach, we identified and merged lexical variants (e.g., “covid19” and “covid-19”, “neurodegenerative” and “neurodegenerative disease”) that would otherwise fragment network centrality and path counts. This normalization had minimal impact on highly curated sources and the text mining tool that applied standardization of nodes in its pipeline, emphasizing the importance of controlled vocabulary alignment when integrating multi-source biomedical sources, especially for downstream tasks that rely on centrality, connectivity, or shortest-path inference.

Beyond semantic alignment, rigorous quality control of phenotypic associations was essential to ensure biological validity. Our semantic validation pipeline uncovered a key limitation in string-based phenotype matching methods, particularly when aligning terms to phenotype ontologies such as the Human Phenotype Ontology (HPO): a high rate of false positives that risked obscuring meaningful mechanistic insights. To mitigate this, we applied a BioBERT-based semantic analysis to evaluate the contextual plausibility of phenotype–disease associations. This approach effectively filtered out spurious links, retaining only those supported by semantically coherent relationships. Among the validated associations, a statistically significant axis connecting COVID-19, diabetes, and AD emerged—highlighting a shared inflammatory and metabolic signature consistent with recent clinical and mechanistic studies [89], [90] . This finding exemplifies the potential of our approach to generate computationally derived hypotheses with translational relevance.

Future efforts will benefit from advanced harmonization strategies, such as ontology-driven mappings and embedding-based semantic alignment, which can further reconcile inconsistencies and enhance data interoperability. While our initial string-based matching introduced noise, embedding-based validation revealed a subset of semantically coherent, recurrent phenotypes (e.g., neuroinflammation, memory loss) across diverse KG paths. These recurring patterns, although exploratory, were statistically supported via permutation testing and merit further investigation as hypothesis-generating signals.

Finally, we acknowledge several limitations related to data extraction and currency. API-based access, though efficient, may omit visual or contextual information available through KG-specific user interfaces, potentially excluding key mechanistic insights. Additionally, some KGs—such as SCAI-DMaps—have not been recently updated and may lack emerging knowledge on COVID-19-related neurological complications. Ensuring the relevance and accuracy of KGs requires ongoing investment in manual curation and system maintenance, which remains resource-intensive but essential for sustaining analytical validity over time.

### 4.7 Text Mining Limitations and Future Directions

Our systematic error analysis of the Sherpa text mining pipeline revealed several critical limitations that likely affected both the completeness and accuracy of the extracted mechanistic relationships. A key challenge was Sherpa’s frequent misinterpretation of speculative language and hedging statements as definitive causal associations. Phrases such as *“it is still debated whether”* were often treated as established facts rather than signals of uncertainty. This issue was especially pronounced in the COVID-19 neurological literature, where much of the mechanistic understanding remains provisional and rapidly evolving. As a result, some false positive associations may have been introduced into our hypothesis database, potentially inflating the perceived certainty of certain biological relationships.

In addition, Sherpa demonstrated difficulties in capturing complex causal chains accurately, often collapsing multi-step biological pathways into oversimplified direct links. For example, while direct relationships such as TLR4 activation leading to TNF-α expression were identified, intermediate steps—such as NF-κB upregulation—were frequently overlooked. This simplification risks obscuring important mechanistic details and distorting the true sequence of biological events relevant to COVID-19-neurodegeneration crosstalk.

Furthermore, our analysis highlighted a tendency for Sherpa to under-extract relationships in sentences containing multiple mechanistic links. Typically, only the first relationship mentioned was captured, with secondary associations systematically missed. This under-extraction suggests that the mechanistic overlap we report likely represents a conservative estimate, emphasizing the most robust and frequently discussed pathways in the literature, while more nuanced, context-dependent, or speculative relationships remain under-represented.

To overcome these limitations, future pipeline improvements should leverage advances in NLP techniques. The use of contextual embeddings and large language models (LLMs) holds promise for more accurately detecting negations and speculative language, thereby reducing false positives arising from uncertainty in the text. LLMs also offer the potential to better capture complex, multi-step causal chains and to recognize secondary, context-dependent associations that current rule-based systems often miss. Coupling LLMs with retrieval-augmented generation (RAG) frameworks could further enhance performance by grounding model predictions in up-to-date, domain-specific knowledge bases, improving factual accuracy and reducing hallucinations. Such an approach would enable the pipeline to dynamically retrieve relevant background knowledge when interpreting subtle or ambiguous mechanistic claims.

Developing domain-specific entity recognition models tailored specifically to COVID-19 and NDD terminology will also be essential for improving entity identification and ontology mapping accuracy. Complementing these models with uncertainty quantification methods would enable the pipeline to flag low-confidence extractions, helping prioritize relationships for further validation. The integration of rule-based filters, contextual language models, and ontological grounding through resources such as UMLS or MONDO can enhance disambiguation of terms and improve recognition of negated or speculative claims. Restricting extracted relationships to those with high confidence, such as well-established causal mechanisms, may further improve the quality and interpretability of the resulting KGs.

Finally, expanding Sherpa’s scope beyond molecular entities to encompass clinical features, environmental exposures, and behavioral symptoms—potentially through integration with public health and electronic medical records—would enable a more holistic representation of COVID-19 and neurodegenerative disease comorbidities. This comprehensive approach could better capture the multifactorial nature of these overlapping disease processes and inform translational research.

### 4.8 Integrating Demographic and Comorbidity Context into Graph-Based Analysis

While our graph-based analysis uncovers biologically meaningful mechanisms and factors linking COVID-19 and NDDs, an important limitation is that several critical clinical covariates—particularly age, sex, and comorbidity burden—are not yet integrated into the KG structure. This limitation constrains the clinical interpretability and stratification of our hypotheses, as these variables are known to shape both COVID-19 outcomes and NDD trajectories.

The KGs employed in this study—such as DisGeNET, DrugBank, and OpenTargets— primarily capture molecular relationships derived from curated literature, experimental data, and ontologies. While these resources excel at representing gene–disease, protein–pathway, and drug–target interactions, they are largely agnostic to the demographic and clinical contexts in which these relationships are observed. For example, DisGeNET provides robust gene–disease associations but does not stratify these by age, sex, or comorbidity status, despite evidence that genetic susceptibility patterns differ across such subgroups [28] .Likewise, canonical pathway databases (e.g., KEGG) do not account for age-related immune changes or sex-specific neuroinflammatory processes [77], [78].

Demographic variables critically modulate COVID-19 severity and NDD risk. Advanced age is the strongest risk factor for both severe COVID-19 and neurodegenerative decline, mediated by mechanisms such as immunosenescence and chronic inflammation [17]. Sex differences also shape disease manifestations: males show higher acute COVID-19 mortality and neuroinflammatory response, while females are more likely to develop long COVID neurological symptoms or faster cognitive decline in Alzheimer’s disease [1]. Comorbid conditions such as hypertension, diabetes, and cardiovascular disease further interact with these factors to influence outcomes. The current KG design, however, generates hypotheses such as “TLR7-mediated immune dysregulation links COVID-19 to Alzheimer’s disease” without distinguishing whether this mechanism is more relevant, for instance, in elderly males or females with autoimmune predisposition.

The lack of demographic and comorbidity stratification in our models has important implications for translational utility. Therapeutic targets identified through our analyses may exhibit differential efficacy across demographic groups. For example, TRPV4—a calcium channel implicated in our findings—has age-dependent expression patterns that could affect its relevance as a therapeutic target in elderly versus younger populations. Similarly, genetic markers identified (e.g., rs5117-C, rs13107325-T) would require calibration in risk models to ensure predictive accuracy across diverse patient groups. This underscores the need to complement molecular discovery with population-level demographic information to better inform personalized medicine strategies.

Future work should therefore aim to systematically integrate such demographic and clinical covariates into graph-based comorbidity analyses. In this regard, several approaches are promising:

- Node attributes: Annotating nodes with metadata (e.g., age group, sex, comorbidity index) to enable subgroup filtering.
- Edge qualifiers: Weighting edges based on subgroup-specific evidence, such as sex-stratified GWAS or age-specific transcriptomic patterns.
- Graph filtering and stratified subgraphs: Extracting hypotheses specific to subpopulations, such as elderly patients with cardiovascular comorbidities.

Resources like UK Biobank and N3C [91] offer large-scale, richly annotated EHR and genomic datasets that could facilitate such integration. Methodologically, advanced network representations—such as multiplex or layered graphs—can explicitly model heterogeneity, while graph neural networks may enable learning embeddings that reflect both molecular and demographic dimensions [85].

Despite these limitations, our findings highlight fundamental mechanisms—particularly immune and inflammatory pathways—that are likely relevant across demographic groups, albeit with varying intensities. These provide a strong foundation for hypothesis-driven experimental validation that can further refine demographic specificity. We would consider that future research efforts prioritize:

1. Collaboration with longitudinal clinical cohorts maintaining demographic and comorbidity data.
2. Development of graph construction pipelines that preserve stratification information from source datasets.
3. Sensitivity analyses to evaluate the robustness of findings across subgroups.
4. Validation of mechanisms in demographically diverse experimental models.
5. Standardization of methods for representing and querying demographic data in biomedical KGs.

The absence of demographic covariates reflects a broader challenge in comorbidity modelling: translating molecular-level insights into patient-level, personalized predictions. While our study aimed at advancing understanding of COVID-19–NDD comorbid mechanisms, achieving true clinical utility will require models that reflect the heterogeneity of real-world populations. By integrating molecular, demographic, and clinical data, future graph-based frameworks can deliver more precise, equitable, and actionable hypotheses that support both scientific discovery and personalized care.

### 4.9 From Comorbidity Hypothesis Generation to Causal Modeling: Advancing Knowledge-Based Discovery

While our current approach successfully identifies associations and comorbidity patterns between COVID-19 and NDDs across multiple KGs, a limitation remains: distinguishing between mere correlations and genuine causal relationships. Traditional graph-based analyses, including our shortest path analysis and phenotype coverage exploration, primarily capture associative patterns without establishing causal directionality or mechanistic causality. This limitation is particularly critical in the context of COVID-19 and NDDs, where understanding whether SARS-CoV-2 infection directly triggers neurodegenerative processes, accelerates existing pathology, or simply co-occurs with neurological symptoms has profound implications for prevention and treatment strategies.

Recent advances in knowledge-based causal discovery offer promising solutions to this challenge. Unlike statistical causal discovery methods that rely on observational data, knowledge-based approaches leverage the semantic content and mechanistic information encoded within KGs to infer causal relationships between variable pairs [92]. This methodology is particularly well-suited to our multi-scale graph analysis framework, as it can operate on the rich biological context present in resources like PrimeKG, SCAI-DMaps, and our curated databases.

The integration of knowledge-based causal discovery into our framework would involve three key enhancements. First, metapath-based causal ranking could replace our current connectivity-focused path analysis. Instead of simply counting paths between COVID-19 and NDD entities, we could identify causally informative metapaths—such as [virus → inflammatory mediator → blood-brain barrier disruption → neurodegeneration]—and rank them based on their causal relevance. Second, LLM-guided causal inference could be applied to our identified pathways, leveraging the extensive biological knowledge embedded in large language models to distinguish between correlation and causation. For instance, the pathway connecting COVID-19 to neuroinflammation via cytokine storm could be evaluated not just for connectivity, but for the temporal sequence, biological plausibility, and mechanistic sufficiency of the proposed causal relationship, using the supporting evidence. Third, causal confidence scoring could be integrated into our hypothesis database, providing each proposed mechanism with a quantitative measure of causal evidence strength.

Such an approach would be particularly valuable for interpreting our findings regarding shared genetic markers like TLR7, VEGFA, and APOE. Rather than simply noting their association with both COVID-19 severity and neurodegeneration, a causal discovery framework could help determine whether these represent: (1) causal pathways where COVID-19 triggers neurodegeneration through these molecular mechanisms, (2) shared vulnerability factors that independently contribute to both conditions, or (3) downstream consequences of a common upstream cause. This distinction is crucial for therapeutic targeting and risk stratification.

Furthermore, the temporal dimension inherent in causal relationships could help address one of the key questions in COVID-19 research: the relationship between acute infection, long COVID, and long-term neurological complications. Our current analysis captures associations between COVID-19 and neurodegenerative pathways but cannot distinguish between immediate inflammatory responses that resolve without lasting damage versus persistent mechanisms that drive progressive neurodegeneration.

The implementation of knowledge-based causal discovery would also enhance the clinical utility of our hypothesis database. Instead of presenting researchers with lists of associated pathways requiring experimental validation to determine causality, we could provide causally-ranked mechanisms with confidence scores, enabling more targeted and efficient experimental design. This is particularly important given the resource constraints in translating computational findings to therapeutic applications.

### 4.10 From In-Silico Prediction to Experimental Validation: Future Directions

Finally, while this study presents a systematic computational framework for uncovering and harvesting potential mechanisms and hypotheses linking COVID-19 and neurodegenerative disorders, it is important to emphasize that all conclusions currently rest on in-silico analyses. These hypotheses, although grounded in high-quality curated knowledge and rigorous network-based reasoning, remain unvalidated by direct experimental, imaging, or cohort-level evidence. Therefore, experimental confirmation is a critical next step to ensure the robustness, biological plausibility, and translational relevance of these proposed mechanisms.

Several complementary strategies can be employed to bridge this gap. Computational validation techniques—such as reverse causal reasoning, network perturbation analyses, candidate mechanism amplitude scoring, and patient-level clinical embeddings [93], [94], [95]—provide a valuable first layer of plausibility assessment. These tools help prioritize mechanisms by integrating evidence from diverse datasets and quantifying their expected impact. However, computational approaches alone cannot replace direct biological testing.

Accordingly, wet-lab and clinical validation through molecular, cellular, and imaging-based assays are essential. Laboratory experiments can test predicted gene–phenotype or pathway– disease associations in relevant models, such as neuronal cell cultures, brain organoids, or animal models. Similarly, clinical studies—leveraging patient registries, neuroimaging data, and longitudinal cohorts—can examine whether the hypothesized mechanisms manifest in real-world patient populations, particularly those with long COVID or NDD exacerbations. We recognize that experimental validation of all possible hypotheses is infeasible due to resource constraints, and therefore, our computational prioritization aims to highlight the most promising targets for focused validation.

Yet, standardized validation protocols, evidence grading, and automated workflows for data integration help maintain rigor and transparency in this regard. The COMMUTE consortium—comprising computational biologists, clinical neurologists, infectious disease experts, and wet-lab scientists—provides an institutional framework to coordinate and accelerate this validation effort across Europe.

Furthermore, the comorbidity database supports integration into biomedical research pipelines. Clinicians and translational scientists can use it to explore ranked hypotheses for biomarker discovery, therapeutic target identification, and drug repurposing efforts, informed by both computational predictions and accumulating experimental evidence. Early adopters within the research community have already begun to validate some of the predicted factors and mechanisms [96], [97], demonstrating the utility of this framework as a starting point for hypothesis-driven experimentation.

Finally, given the rapidly evolving landscape of COVID-19 research and the continuous generation of new biomedical data, it is imperative that the co-morbidity graphs and hypothesis rankings remain up-to-date. Automated and adaptive update mechanisms ensure that our framework remains relevant and accurate, reflecting the latest scientific advances.

By explicitly acknowledging the current limitations of computational inference and outlining a clear path toward systematic experimental validation, this study lays a foundation for translating in-silico insights into actionable knowledge. Future efforts to integrate diverse experimental and clinical datasets with computational models will be pivotal in advancing our understanding of COVID-19–related neurological complications and informing the development of targeted therapeutic interventions.

## 5 Conclusion

This study utilized a diverse collection of databases and resources, structured as KGs, alongside advanced text mining techniques, to investigate co-morbidity mechanisms between COVID-19 and NDDs. Through graph-based analyses and phenotype coverage explorations, we highlighted how diverse KGs contribute unique insights into the molecular and clinical intersections between these conditions. Smaller KGs offer targeted pharmacological data, while larger and medium-sized KGs capture broader systemic interactions, particularly in inflammation and immune responses. The study underscores the need for an integrated approach to capture these multifaceted connections but also recognizes limitations such as data heterogeneity, the need for continuous updates of KGs, and the lack of experimental validation. Future enhancements — including advancements in text mining tools and the integration of diverse data types such as clinical records, electronic health records (EHRs), and real-world evidence — hold significant potential to deepen our understanding of comorbid pathways. By bridging molecular insights with clinical observations, these advancements could provide a more holistic view of disease mechanisms and facilitate the identification of actionable therapeutic targets.

Beyond elucidating disease mechanisms, our publicly available co-morbidity hypothesis database provides a versatile resource for advancing both research and practice. By systematically mapping molecular and clinical intersections between COVID-19 and NDDs, it supports hypothesis generation, patient risk stratification, and identification of druggable pathways. Its accessibility enables collaborative validation and continuous refinement, fostering translational research and data-driven therapeutic strategies. Overall, our findings underscore the promise of graph-based approaches to not only deepen mechanistic understanding but also guide targeted interventions and improve outcomes for vulnerable populations.

## Data and Code Availability

The data and source codes used in this study are available at: https://github.com/SCAI-BIO/covid-NDD-comorbidity-NLP.

## Acknowledgement

The authors would like to thank Prof. Laura Furlong, IMIM Barcelona, for her support with the DisGeNet analysis. The authors would also like to thank Dr. Benjamin Gyori, Harvard Medical School, for fruitful discussions and guidance with the INDRA system.

